# Hierarchical Assembly of Single-Stranded RNA

**DOI:** 10.1101/2023.08.01.551474

**Authors:** Lisa M. Pietrek, Lukas S. Stelzl, Gerhard Hummer

**Affiliations:** Department of Theoretical Biophysics, Max Planck Institute of Biophysics, Max-von-Laue-Straße 3, 60438 Frankfurt am Main, Germany; Faculty of Biology, Johannes Gutenberg University Mainz, Gresemundweg 2, 55128 Mainz, Germany; KOMET 1, Institute of Physics, Johannes Gutenberg University Mainz, 55099 Mainz, Germany; 4Institute of Molecular Biology (IMB), 55128 Mainz, Germany; Institute for Biophysics, Goethe University, Max-von-Laue-Straße 9, 60438 Frankfurt am Main, Germany

## Abstract

Single-stranded RNA (ssRNA) plays a major role in the flow of genetic information– most notably in the form of messenger RNA (mRNA)–and in the regulation of biological processes. The highly dynamic nature of chains of unpaired nucleobases challenges structural characterizations of ssRNA by experiments or molecular dynamics (MD) simulations alike. Here we use hierarchical chain growth (HCG) to construct ensembles of ssRNA chains. HCG assembles the structures of protein and nucleic acid chains from fragment libraries created by MD simulations. Applied to homo- and heteropolymeric ssRNAs of different lengths, we find that HCG produces structural ensembles that overall are in good agreement with diverse experiments including nuclear magnetic resonance (NMR), small-angle X-ray scattering (SAXS), and single-molecule Förster resonance energy transfer (FRET). The agreement can be further improved by ensemble refinement using Bayesian inference of ensembles (BioEn). HCG can also be used to assemble RNA structures that combine base-paired and unpaired regions, as illustrated for the 5^1^ untranslated region (UTR) of SARS-CoV-2 mRNA.

## 1 INTRODUCTION

Single-stranded RNAs (ssRNAs) play important roles in many cellular processes, in particular in the transmission of genetic information in the form of messenger RNA (mRNA). Non-coding stretches in mRNA or fully noncoding ssRNAs have key roles in the regulation of transcription and translation, e.g., by acting as riboswitches^1^ or by regulating the nuclear export of mRNA, its stability and translation via polyadenylation.^2,3^ In solution ssRNAs can remain dynamically fully flexible and unstructured, transiently adopt secondary structures with paired bases, or form more permanent secondary structure in complex with a binding partner.^4^ In recent years, ssRNA-based therapeutics have come into focus for a variety of applications. Small molecules binding to aptamers and thus affecting gene expression have been investigated in disease treatment such as cancer or bacterial infections.^5^ Modulating expression by increasing gene accessibility or by gene silencing is another exciting prospect in the field of microRNA research. ^6^ Therapeutics based on mRNA have long been subject for discussion.^7^ Recent advances in RNA research cleared the path for mRNA therapeutics,^8^ such as vaccines based on mRNA.^9^

In order to improve our understanding of ssRNA and their functional mechanisms we need to characterize their structural and dynamical features. However, experimentally investigating disordered ssRNA remains a challenging task. In recent years, nuclear magnetic resonance (NMR) techniques have been proven powerful to investigate local structure and dynamics with high-resolution in short disordered stretches of ssRNA shifts.^10–15^ Smallangle X-ray scattering (SAXS) studies^16–18^ or Förster resonance energy transfer (FRET) techniques^16,19–21^ yield insight into global structure of flexible biomolecules. The negatively charged ssRNA molecules have been shown to be strongly dependent on environmental buffer conditions, including ion concentration and type,^12,16–18^ an effect seen also in molecular dynamics (MD) simulations.^10,21^

Structural ensembles of ssRNA that capture the heterogeneity of these highly dynamic systems at atomic detail help the interpretation of data from experiments. Most experiments report on ensemble averages. Such ensembles can, in principle, be obtained by performing MD simulations. However, MD simulations suffer from inaccuracies in the available force fields.^22,23^ For RNA, special care has to be taken in setting the buffer conditions and choosing the ion force field parameters.^24,25^ For biopolymers, small systematic errors in, say, backbone torsion potentials add up and result in major structural imbalances.^26^ Inaccuracies in the energetics are amplified by the broad and shallow energy landscape of flexible biomolecules,^13,21,27,28^ which requires extensive sampling. The sampling of ssRNA structural ensembles by MD simulations thus suffers both from systematic uncertainties due inaccuracies in the force field and from statistical uncertainties due to the slow structural dynamics. Fragment assembly is a promising approach to model RNA 3D structures. In early applications of RNA fragment assembly, Das et al. used FARFAR, a Rosetta-like fragment assembly approach to model noncanonical double-stranded (dsRNA) structure with atomistic detail.^29^ Their fragment structures were drawn from a library based on RNA structure in the large ribosomal subunit.^29^ The more recent FARFAR2 approach^30^ has been used to generate ensembles of short ssRNA polymers.^14^ Chojnowski et al. developed a method to model 3D structures of short RNA polymers featuring base-paired strands as well as unpaired strands involved in loops by assembling RNA fragments from the PDB with the option to include experimental restraints.^31^

Ensembles of flexible biopolymers can be improved by integrating experimental data. Approaches such as Bayesian/Maximum Entropy (BME)^32–35^ and Bayesian inference^36–40^ have been shown to work well in applications to ensembles of disordered biomolecules. For instance, Bottaro and co-workers refined tetrameric fragments according to NMR data using a BME approach, to improve their structural ensembles obtained via MD simulation, resulting in a more accurate description of the thermodynamic states. ^13^ In another example, Bergonzo et al. showed that a BME approach helped to improve conformational ensembles of a heteropolymeric oligonucleotide by integrating NMR and SAXS experimental data.^14^ Alternatively, integration of experimental information can help to build models of observed molecules.^18,41,42^

In previous work, we have introduced the hierarchical chain growth (HCG) method for disordered proteins.^28,40,43^ We found that HCG is a robust approach well suited to efficiently grow broad structural ensembles of disordered proteins with atomic detail that are in line with experimental findings. Here, we adapted HCG to model structural ensembles of disordered ssRNA. We focus on systems for which experimental data are available as reference:^14,18,21^ homopolymeric adenosine monophosphate multimers (rA_n_ with *n* = 19, 30), homopolymeric uridine monophosphate 30mer (rU_30_), and the short disordered heteropolymeric ssRNA rUCAAUC. In particular, ssRNA fragment assembly is implemented in the form of a Monte Carlo chain growth algorithm, which we then used to assemble structural ensembles of conformations with atomic detail. We validated the modeled ensembles against diverse experimental data and could establish good agreement on average already without refinement. We further improved the agreement with experimental observations by integrating experimental data using BioEn as a gentle ensemble refinement method.^37^ As a proof of principle, we demonstrate that ssRNA chains grown with HCG can be combined with models of dsRNA, paving the way towards modeling short unstructured linkers, terminal untranslated regions (UTRs) or loops.

## 2 METHODS

### 2.1 MD Fragment Library

For poly-adenine (A) RNA, an rA_4_ tetramer was modeled using the AMBER suite of programs.^44^ The oxygen atoms of the terminal ribose groups at the 5^1^ and 3^1^ ends were protonated (see Figure S1 for rA_4_). For heteropolymeric ssRNA, we used heterotetrameric fragments rGXYZ. The nucleotide at the 5^1^ position was fixed as guanine (G). We chose guanine as head group, first, to mimic the interior of the ssRNA by providing a purine platform for stacking and, second, to facilitate the alignment with a relatively large base. For the following three nucleotides ‘XYZ’, we used all 4^3^ = 64 combinations of G, A, cytosine (C), and uracil (U). Each RNA fragment was placed in a dodecahedral box and solvated in TIP4P-D water^45^ with 150 mM NaCl.^46^ Charge neutrality was established with excess sodium ions. On average, resulting systems contained about 6600 atoms in total. The RNA fragments were modeled with the DESRES^23^ force field. We thus performed the fragment MD simulations using the same force field, water model and ion parameters as described before.^21^

MD simulations were performed with GROMACS/2018.8.^47^ Bonds including hydrogen atoms were constrained using the P-LINCS algorithm.^48^ To maintain the pressure at a constant value of 1 bar, the Parrinello-Rahman barostat^49^ was used. The cut-off distances for van der Waals and electrostatic interactions were set to 1.2 nm. Electrostatic interactions were calculated using the particle mesh Ewald method^50^ with the Fourier spacing set to 0.16 nm. The system was first energy minimized, followed with 400 ps of MD equilibration. The production REMD simulation was run in the NPT ensemble for 100 ns with structures saved every 10 ps. For all tetramer fragments we used 25 replicas that collectively spanned a temperature range of 300-431 K, as calculated using the algorithm by Patriksson and van der Spoel.^51^ For each system, 10000 different structures collected at equally spaced time points from the replica simulated at 300 K were used for the respective fragment library.

### 2.2 Hierarchical Chain Growth

We adapted HCG^40,43^ to grow full-length models of disordered homo- and heteropolymeric ssRNA chains from MD rA_4_ and rGXYZ fragments. HCG was previously implemented and validated to model extensive ensembles of intrinsically disordered proteins (IDPs), displaying average properties that are in line with experimental observables.^28,40,43^ HCG performs fragment assembly, i.e., a pool of fragment structures are then combined at random into long polymers. The structural alignment of individual fragments and the rejection of poorly aligned or sterically clashing fragment pairs are critical for the quality of the resulting ensembles in terms of both local and global structural properties. We note that besides the root-mean-square distance (RMSD) alignment criterion and the steric exclusion we did not include any kind of attractive or repulsive inter-fragment interaction during the assembly. Thus, the electrostatic interaction considered here is intra-fragment interactions sampled in the fragment MD simulations. However, we integrate experimental data on a global level to account for resulting discrepancies in assembled polymers.

We used a fragment alignment strategy that focuses on the nucleic acid backbone but also accounts for the position of the base. For two conformations of fragments adjacent in sequence and drawn at random from the respective pool, we performed a rigid-body superimposition of the O3^1^ atom, phosphate atom, and O5^1^ atom connecting nucleotides *−*2 and *−*1 in fragment 1 and nucleotides 1 and 2 in fragment 2, and all atoms of the nucleobase as well as the C1 atom of nucleotide *−*1 in fragment 1 and of nucleotide 2 in fragment 2 (Figure 1, light blue shaded area). For a successful alignment, we required the RMSD of the superimposed atoms to below a given threshold, RMSD *<* 0.64 Å. In the superimposition, we doubled the weights of the aligned backbones atoms relative to the aligned atoms of the nucleobase to produce atom distances within the expected range of experimentally determined covalent radii of atoms.^52^

**Figure 1:**
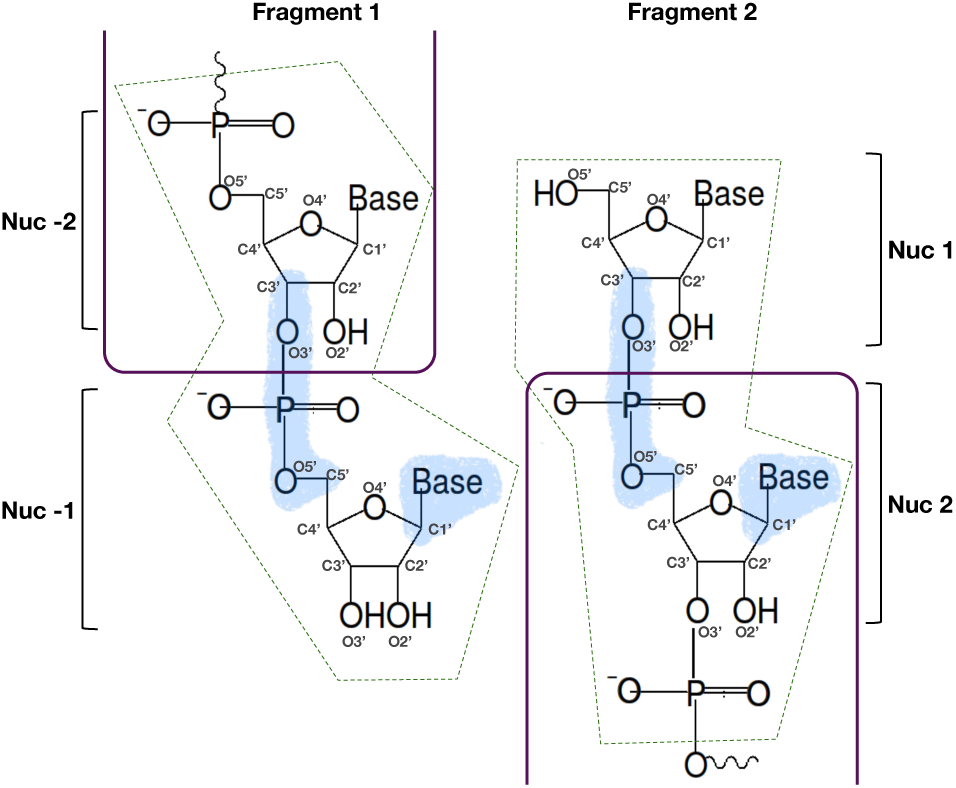
Fragment alignment. Aligned heavy atoms are highlighted by blue shade. Heavy atoms within the dashed dark green box were excluded from the clash search. The purple lines indicate the regions of the two fragments that are included in the assembled chain.

Alignment was followed by a search for steric clashes, defined as heavy atom distances below a cut-off of 2 Å. Note that we did not consider hydrogen atoms in clash detection. Atoms in the fragment-overlap region were excluded from the heavy atom clash search (Figure 1, dark green dashed boxes). Any steric clash resulted in the rejection of the pair. Otherwise, the two fragments were merged. In merged fragments, nucleotide *−*1 from the first and nucleotide 1 from the second fragment were removed. In this way, the assembled chain featured only nucleotides sampled at the second and third position (X, Y) of the rGXYZ fragments. The terminal nucleotides (G, Z) were treated as capping groups. We repeated this procedure in each hierarchical level of HCG until we reached the full-length sequence. For each polymer investigated in the present work we grew ensembles with 10000 members. The RNA structure libraries (i.e., the MD fragment library as well as exemplary structures from the HCG ensembles discussed in this work) are available at https://zenodo.org/record/8369324. The HCG code to assemble ssRNA to the hierarchical chain growth is available at the GitHub repository https://github.com/bio-phys/hierarchical-chain-growth/.

We note that the ribose atoms are not included in the superimposition. We found that by not enforcing the sugar pucker configuration we increased the diversity of grown structures and benefit from diverse sugar pucker configuration sampled in fragment MD simulations. By including the nucleobase in the alignment, we improved the configuration of stacked bases, which is important to produce reasonable stacking also for longer sequences (Figure 2). The extent of base stacking in the assembled structures will to a significant degree be predetermined by the fragment library entering HCG, and thus the MD simulation force field used to create the library.^12^

**Figure 2:**
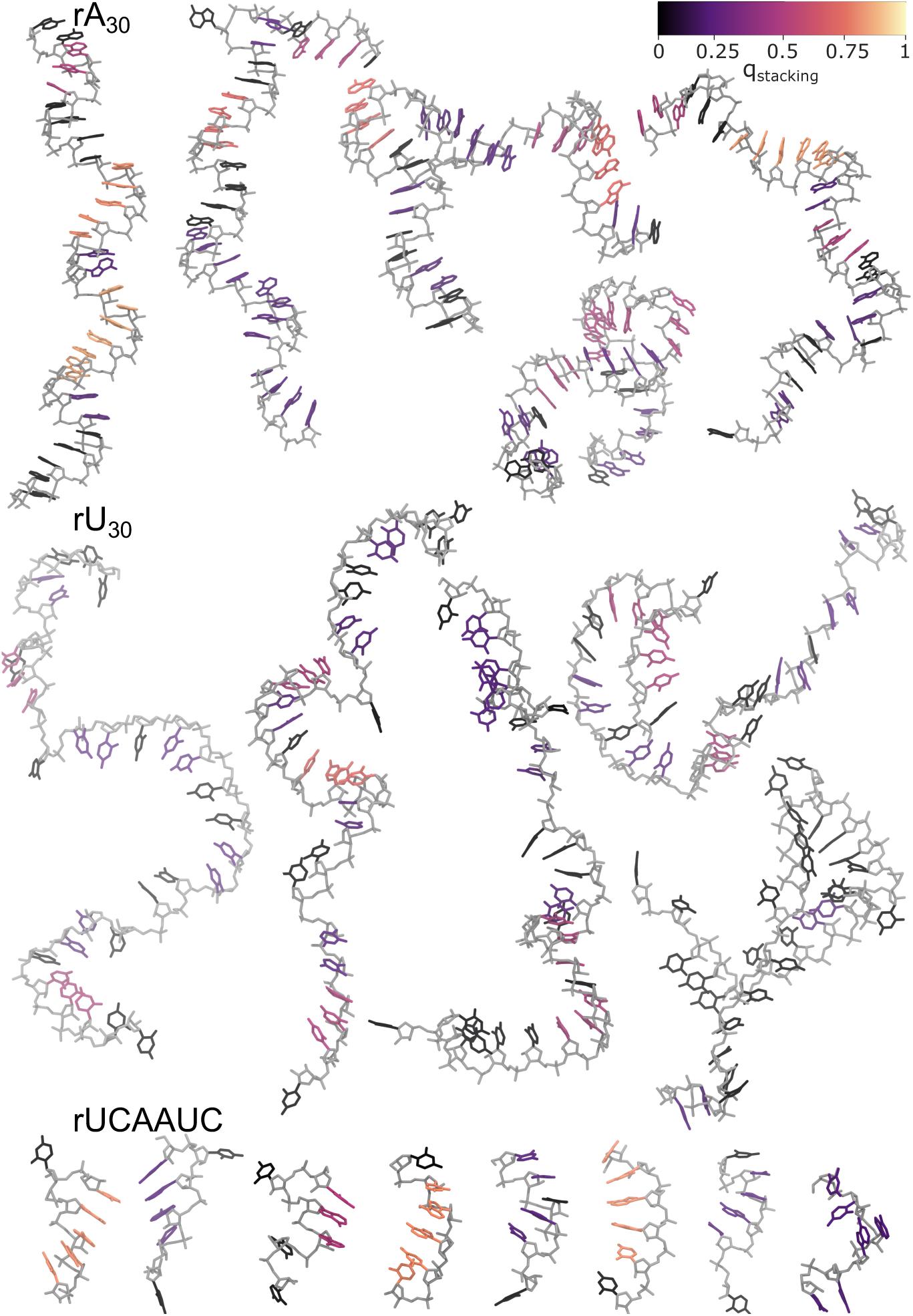
Snapshots of ssRNA polymers grown with HCG. Representative renders of structures drawn at random from ensembles of (top) rA_30_, (center) rU_30_, and (bottom) rUCAAUC sampled by fragment assembly. The nucleic backbone is shown in light gray and nucleobases are colored according to the stacking. Hydrogen atoms are omitted for clarity.

### 2.3 Modeling the 5***^t^*** UTR of SARS-CoV-2 mRNA

To build a structural ensemble of the 5^1^ UTR of SARS-CoV-2 mRNA by HCG, we combined fragments for the ssRNA segments with structural models for the stem loops using the secondary structure as input. The conformations of the structured stem-loop regions were randomly drawn from libraries filled with MD trajectories published previously. ^53^ The connected disordered regions were grown using HCG as described above according for the sequence in Ref.^53^ For the assembly the same scheme as implemented in HCG was used. In particular, the adjacent regions (fragments) were assembled in a hierarchical manner in subsequent levels. In the final level, the full-length model was assembled with a total of 233 nucleotides (sequence and secondary structure in Supplementary Figure S2). For the heavy atom superimposition we set the RMSD cut-off to 1 Å, the clash radius was kept at 2 Å. We grew only a small ensemble with 50 full-length chains. We note that the region spanned by nucleotides 162-200 was predicted to be structured and a part of stem-loop SL5.^54^ However, to our knowledge for this region there has been no structure solved so far. Therefore we here modeled this region as single-stranded region with HCG.

### 2.4 Mapping of FRET Labels

HCG is naturally suited to the inclusion of molecular labels such as covalently attached fluorophores. To build a pool of dye-labeled rA_4_ fragments, we used an MD library for the dyes Alexa Fluor 594 and Alexa Fluor 488 attached to tetradeoxyadenosinemonophosphate (dA_4_). The use of dA_4_-dye fragments to model fluorophores attached to both DNA and RNA chains has been validated by Grotz et al.^21^ A random structure was drawn from the pool of rA_4_ fragments and from the Alexa 594 or Alexa 488 MD library, to either label the 5^1^ or 3^1^ end, respectively, of the rA_4_ fragments. We performed a rigid body alignment of heavy atoms of the sugar moiety and nucleobase from the terminal nucleotides. In particular, to attach Alexa Fluor 594 to rA_4_ we aligned the respective atoms from the terminal nucleotide at the 5^1^ end of the rA_4_ fragment with the respective atoms from the terminal nucleotide at the 3^1^ end from the dA_4_-dye fragment. For Alexa Fluor 488 the same alignment was performed but at the 3^1^ end of the rA_4_ fragment and at the 5^1^ end from the dA_4_-dye fragment. The RMSD cut-off for heavy atom distances was set to 0.8 Å. If the RMSD value was below the cut-off, we searched for clashing heavy atoms within a pair distance of 2.0 Å. If no clashing atoms were detected, the dye molecules and the rA_4_ fragment were assembled such that all atoms from the dA_4_ fragment and terminal oxygens of rA_4_ were excluded. In this way we sampled a library of the FRET dyes mapped onto rA_4_ fragments with 10000 conformations for both fluorophores each, Alexa Fluor 594 and Alexa Fluor 488.

### 2.5 Ensemble Reweighting Using BioEn

We refined the HCG ensembles of the ssRNA polymers investigated here against experimental SAXS or single-molecule FRET data by reweighting using the Bayesian Inference of Ensembles (BioEn) procedure.^37,38^ We used uniform reference weights *w*_0_*_,i_* = const. for the unbiased ensembles produced by HCG. The reference weights of the individual chains then were minimally adjusted such that the ensemble average better agrees with the experimental observable, while making sure that the refined ensemble was well-defined and converged. The confidence parameter *θ* in BioEn^37,38^ was chosen by L-curve analysis. As a measure of the extent of reweighting, we used the Kullback-Leibler divergence *S*_KL_ = Σ*_i_ w_i_* ln(*w_i_/w*_0_*_,i_*) between the original weights *w*_0_*_,i_* and the refined weights *w_i_*, both normalized, Σ*_i_ w*_0_*_,i_* = Σ*_i_ w_i_* = 1. In addition, we inspected the cumulative distribution function (CDF) of the rank-ordered weights *w_i_*. A rapid initial rise indicates that few ensemble members carry a large fraction of the weight, which in turn indicates poor overlap between reference and refined ensembles.

### 2.6 Analysis of ssRNA Conformations

The python packages Barnaba,^55^ MDTraj,^56^ and MDAnalysis^57,58^ python libraries were used to perform analyses of the ssRNA conformations.

A cluster analysis was performed using the Barnaba software^55^ exemplary of the rA_4_ fragment conformations as sampled in 100 ns MD simulation trajectory. The sampled conformations within 10000 frames were assigned to 6 different clusters. In the cluster analysis, we first calculated the g-vectors describing the relative position of nucleotide pairs in each structure. We then performed a principal component analysis (PCA) of the g-vectors, projecting the data points onto the plane of the first and second principal component axis. The clustering was performed via a Barnaba wrapper of the DBSCAN function from the scikitlearn package. Here, we set the minimum distance for nearest neighbors to eps = 0.35 and the minimum number of samples per cluster to 50.

#### Quantification of Base Stacking

We used the Barnaba software^55^ to screen the assembled structures for stacked bases. For each base, we quantified the stacking by a factor *q*_stacking_. For nucleobases not involved in any stack, we set *q*_stacking_ = 0. For stacks of *n*_stacked_ = 2 we set *q*_stacking_ = 0.25*c* and *q*_stacking_ = 0.5*c*, respectively. For longer consecutive stacks with *n*_stacked_ *≥* 4 bases, we set *q*_stacking_ = *c − c/*(*n*_stacked_ *−* 1). We set *c* = *n/*(*n −* 1), where *n* is the number of nucleobases, so that in a complete stack *q*_stacking_ = 1 for all bases. We then used *q*_stacking_ to color the bases in structural visualizations (Figure 2).

### 2.7 Calculation of Experimental Observables

#### FRET

We calculated FRET efficiencies for the rA_19_ ssRNA ensembles obtained by HCG with explicit dyes attached at the 5^1^ and 3^1^ ends. The inter-dye distance *r* was calculated as the geometric distance between the central oxygen atoms of the two FRET dye labels, ^21^ as determined using MDAnalysis.^57,58^ For the orientational factor *κ*^2^ in the Förster theory we considered three models that differed in their assumptions on the dye dynamics. A similar approach has been employed before.^59^

In model 1, we set *κ*^2^ = 2*/*3,^40,60^ assuming implicitly that dye rotation is isotropic and fast^61,62^ compared to the fluorescence lifetime of the donor, which in absence of the acceptor is *τ_D_ ≈* 4 ns.^21^ The transfer efficiency *E* of an individual ssRNA conformation labeled with fluorophores at each end was then calculated as

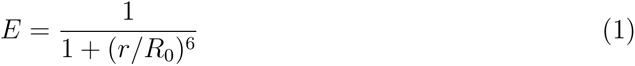

The Förster radius *R*_0_ was set to the experimentally determined value of *R*_0_ = 5.4 nm.^21^ In model 2, we assumed also the dye linker dynamics to be fast and accordingly attached *≈*20 dye pairs to a given ssRNA conformation, averaged the inter-dye distance *r* over these conformers, and then calculated the FRET intensity according to eq. 1 with the average *r*. By contrast, in model 3 we assumed the dye dynamics to be slow. Accordingly, we determined both *r* and *κ*^2^ explicitly for each dye-labeled ensemble member. We calculated *κ*^2^ as

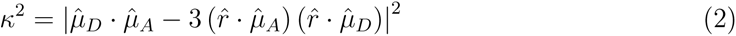

where *µ̂_D_* and *µ̂_A_* are unit vectors in the direction of the transition dipole moments of donor and acceptor, respectively, and *r̂* is a unit vector pointing in the direction between the central oxygen atoms of the two dyes. We then calculated the rate of energy transfer as *k_T_* = (3*/*2) *κ*^2^ *k_D_* (*R*_0_*/r*)^6^ and the FRET efficiency of each ssRNA conformation in the ensemble as^63^

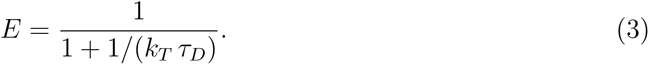

For all three models, the efficiency *E* was then averaged over the ssRNA conformations in the ensemble with their respective weights. The three models can be thought of as extremes with respect to the assumed dye dynamics. Importantly, in all models we assumed the ssRNA dynamics do be slow compared to the fluorescence lifetime *τ_D_*.

For model 2 we mapped FRET labels onto the full-length rA_19_ grown for analysis with model 1 with labels integrated in the models at the fragment level. Particularly, we randomly picked a rA_19_ conformation *i*, attempted to simultaneously replace both labels with randomly picked label conformations of Alexa 594 *j* and Alexa 488 *k*. Here, we followed the procedure for the heavy atom alignment and clash search as described above for the dye mapping on the fragments. In case of a steric clash or if the RMSD exceeded 0.8 Å in the alignment both dye conformations were discarded and a new pair of dyes was drawn. We attempted to replace the dye conformations 1000 times for each of the 10000 randomly drawn rA_19_ conformations. The normalized acceptance rate for dye replacements determined for conformation *i* was then used as weight for *(r_i_)* for each conformation in the ensemble.

#### SAXS

For each ensemble member *i* of either the rA_30_, rU_30_, or rUCAAUC HCG ensembles, we calculated the SAXS scattering intensity *I_i_*(*q*) at scattering vector *q* using Crysol^64^ following Ref.^14^ The calculated scattering intensities *I_i_*(*q*) with normalized weights *w_i_* (i.e., Σ*_i_ w_i_* = 1) were averaged over the ensemble as *I*_sim_(*q*) = Σ*_i_ w_i_I_i_*(*q*). In the limit of *q →* 0, the Guinier approximation becomes exact, *I_i_*(*q*) *≈ I*_0_*_,i_* exp(*−q*^2^ *R_G,i_*^2^*/*3), where *I*_0_*_,i_* is the intensity at *q* = 0 and *R_G,i_* is the radius of gyration of ensemble member *i*. Accordingly, we calculated *R_G,i_*from the slope of ln *I_i_*(*q*) with respect to *q*^2^ at *q* = 0 as

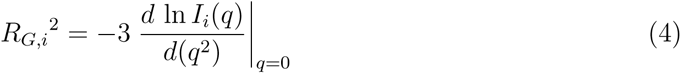

We evaluated the slope as a numerical first difference. With *I_i_*(*q*) *≈ I*_0_*_,i_* exp(*−q*^2^ *R_G,i_*^2^*/*3) at small *q* and *I*_sim_(*q*) = Σ*_i_ w_i_I_i_*(*q*), the apparent *R_G_* averaged over the ensemble is

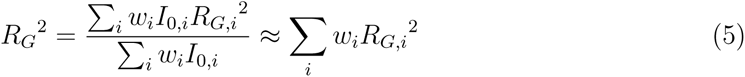

because the *I*_0_*_,i_* calculated by Crysol^64^ are nearly constant, with only small variations caused by differences in the solvent contributions to *I*_0_ across the ensemble. In our analysis, we accordingly report root-mean-square (RMS) *R_G_* values evaluated with ensemble weights *w_i_*. We related the calculated intensity *I*_sim_ and measured intensity *I*(*q*) to each other by performing a least-square fit of *I*(*q*) = *a I*_sim_(*q*) + *b* with an intensity scaling factor *a* and a constant background correction factor *b* as fit parameters. To assess the quality of the fit, we calculated the reduced chi-squared normalized by the number *M* of data points,

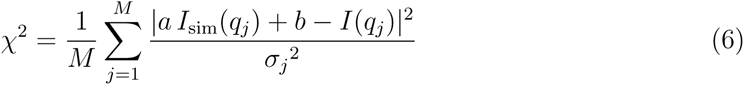

with *σ_j_* the reported experimental standard error of *I*(*q_j_*). Values of *χ*^2^,:S 1 indicate agreement within the experimental uncertainty. We assessed the quality of the ensembles with the *χ*^2^ statistic for the squared residuals and the hplusminus statistic for the sign-order of the residuals, calculating *p*-values for both tests individually and in combination.^65^

## 3 RESULTS AND DISCUSSION

### HCG Produces Broad Structural Ensembles of ssRNA

We used HCG to sample structural ensembles of ssRNA polymers with four different sequences: rA_30_, rA_19_, rU_30_, and rUCAAUC. For all four systems, we observed a combination of extended and compact conformations, as shown representatively in Figures 2 and S3. Compactness is associated with kinks in the ssRNA backbone (see, e.g., bottom center rA_30_ structure in Figure 2). In the more extended structures, the chains retained features of the A-form helix (e.g., rA_30_ top left in Figure 2). In particular, we observed stretches of continuously stacked nucleobases.

The ssRNA structure in the HCG ensembles depends on the nucleotide sequence. Whereas the poly-purine rA_30_ tends to form relatively straight segments of stacked adenines, polypyrimidine rU_30_ is visually rather distorted with stretches of unstacked uridines (Figure 2). A stacking analysis using Barnaba revealed a significant shift from rU_30_ to rA_30_ by about 4 stacks on average (Figure S4A). Here, a single stack was defined as two nucleobases with particular distances and orientation to each other.^55^ We further looked at consecutively stacked nucleobases, which we defined as four or more stacked nucleobases in a row. Again a significant difference in the CDF for both polymers was observed. We colored the nucleobases according to the stacking information from our analysis (Figure 2 and S4B). Key to retaining base stacking in fragment assembly was the inclusion of the atoms of the nucleobase in the RMSD alignment, which ensured that the relative base-base orientation of the fragments was retained in the HCG assembly (Figure 1). The observed sequence dependence is in line with the behaviour previously reported for the ssRNA polymers investigated here.^14,16,18,21,66^ The structure in long ssRNA chains is reflected in the fragments libraries used for HCG. We clustered the rA_4_ fragments library according to their structure. In the largest clusters, stacked A-form like conformations dominate, either with perfectly stacked nucleobases (cluster 0 with 42%) or with nucleotide 4 inverted (cluster 1 with 39%; see Supplementary Figure S1). NMR studies support the presence of a substantial fraction of A-form like conformations for short single stranded RNA fragments.^10,11,67^ The next largest clusters 2 and 3 are sparsely populated (*≈* 1%), containing structures with A3 unstacked and A4 inverted (cluster 2), and all bases unstacked (cluster 3).

HCG assembly largely preserves also the distribution of torsion angles in the fragment libraries, as shown for rA_19_ in Figure S5. In the HCG assembly, the central two nucleotides at positions 2 and 3 of the tetrameric fragments were retained. Figure S5 compares the distributions of the backbone torsions averaged over the 19 bases in each A2 and A3 as sampled in rA_19_ chains to the respective distributions in the fragments. First, we found that the distributions are essentially independent of position in the HCG assembly, as would be expected for a long homopolymer. Second, we found that HCG largely retained the torsion angle distributions of the fragments. However, we observed that some populations were altered or completely vanished. The small differences reflect in part actual incompatibilities with longer chains, yet also choices in HCG, in particular of the atoms to align and of the clash criteria. Differences as, e.g., in *α* for A3 or *E* and *ζ* for both A2 and A3 may be amplified by the different nature of the preceding nucleotide at the 5^1^ position and the following nucleotide at the 3^1^ position of A2 or A3, respectively. In particular, in the MD fragments these nucleotides were attached to terminal nucleotides, which may impact the distribution of torsional angles at the P O5^1^ bond (*α*), C3^1^ - O3^1^ bond (*E*), and O3^1^ - P(+1) bond (*ζ*).

The recovery of the torsional distributions after assembly suggests that as we observed before for tau K18,^40^ the local and global structural features observed in the assembled full-length chain arose from the local structure sampled in the fragments. In particular, rA_4_ fragments sampled in the DESRES force fields mostly exhibited A-form helix like conformation with a considerable population of conformations with one or two of the 4 nucleotides being unstacked (see Figure S1 cluster 1 and 3). In turn this resulted in either pseudo A-form helix-like populations as well as populations of kinked, or loop-like structures for ssRNA polymers sampled with HCG, with some of the chains even featuring patterns with bulging nucleobases (Figure 2 and Figure S3).

### ssRNA from HCG Reproduces SAXS Data

We compared the calculated SAXS intensity profiles obtained by averaging across the rA_30_ and rU_30_ HCG ensembles to experimental profiles measured at 100 mM NaCl^18^ (Figure 3A). The rA_30_ and rU_30_ ensembles were assembled from rA_4_ and rGUUU fragment libraries, respectively (see methods). Overall, the agreement was good, with reduced *χ*^2^ errors (mean-squared residuals divided by the experimental error) of 2.7 and 3.6, respectively. However, the residuals revealed small but systematic deviations for both polymers. In the relatively featureless intensity profiles, the residuals pointed to somewhat too extended structures for rA_30_ and too compact structures for rU_30_.

**Figure 3:**
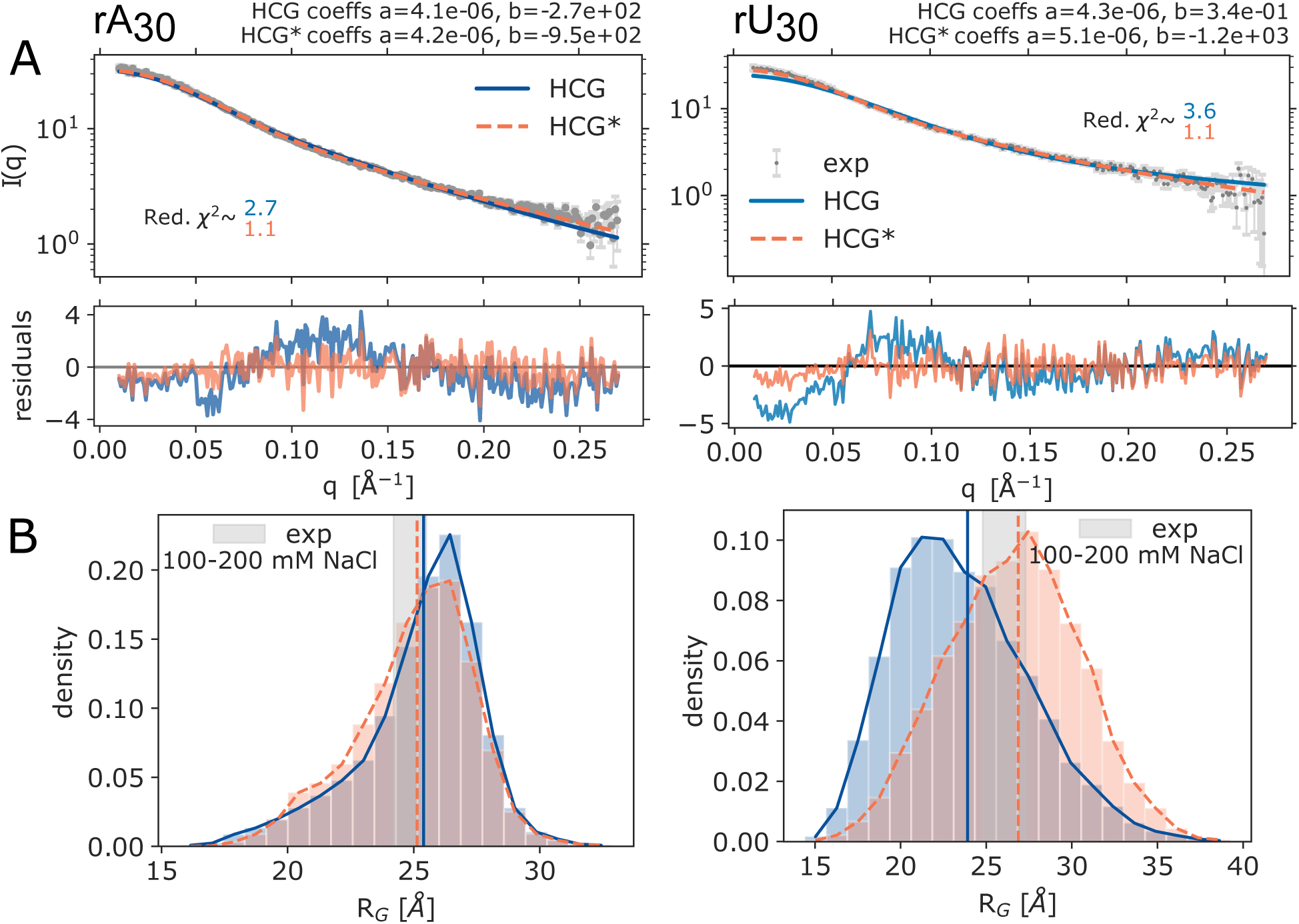
Comparison of the rA_30_ and rU_30_ HCG structural ensemble to SAXS measurements^18^ (left and right column, respectively). (A) Top: Experimental SAXS profiles measured at 100 mM NaCl in gray and the average profile calculated using Crysol^64^ for the unrefined HCG ensembles, 10000 structures each, (blue) and the refined HCG* ensembles with weights for *θ* = 100 (orange). Intensity scale factors *a* and background correction constants *b* determined by least square fitting are shown in the plots. Bottom: Residuals of the experimental and the average profile for the HCG ensembles. (B) Distribution of the *R_G_* in the unrefined HCG as calculated by Crysol and reweighted HCG* ensembles (blue and orange, respectively). Vertical lines indicate the RMS *R_G_* value of the HCG ensemble (solid blue) and the weighted RMS *R_G_* value (HCG*, dotted orange). The gray shaded area highlights the area spanned by the *R_G_* value inferred from the SAXS profiles measured at 100 mM and 200 mM NaCl including the error range.

Considering the fact that HCG does not account for long-range electrostatic interactions and salt screening effects beyond the scale of fragments, the agreement between measured and calculated SAXS intensities at *∼*100 mM NaCl is remarkably good. Solvent and, in particular, ions affect the global structure of the negatively charged nucleic acids polymers.^10,12,16,17,21,24,68^ At the high concentrations of the SAXS experiments, interchain interactions may also be relevant.^18,69^

### Gentle Ensemble Reweighting Further Improves Agreement with SAXS Data

We refined the HCG ensembles of rA_30_ and rU_30_ by performing BioEn reweighting^37,38^ against the experimental scattering profile measured at 100 mM. Using a rather gentle bias we adjusted the weights of the ensemble members to agree with the experimental profile with reduced *χ*^2^ values of *≈*1.1 for both polymers (HCG* ensemble in Figures 3A, S6 light orange, and S7).

We also found good agreement of the measured and calculated values of the radius of gyration, *R_G_*. We calculated the RMS *R_G_* = *(R_G,i_*^2^)^1/2^ as average over the members *i* of the ensemble with their respective weights. For rA_30_, the RMS average *R_G_* over the HCG ensemble was within the uncertainty of the *R_G_* measured by SAXS at 100 and 200 mM NaCl;^18^ for rU_30_, it was just below the expected range (Figure 3B). In this range, salt concentration was found to have only a small effect on *R_G_*.^18,69^ Reweighting by BioEn to match SAXS intensities *I*(*q*) also improved the agreement of calculated and measured *R_G_*values. Overall, HCG captured the global dimensions of rA_30_ and rU_30_, and a gentle BioEn reweighting resulted in near-perfect agreement also at higher NaCl concentrations for rA_30_ (see below).

We tested the impact of the fragment library by comparing the distribution of *R_G_* for rA_3_0 chains grown by HCG with rA_4_ fragments and with rGAAA fragments. We found that the *R_G_* distributions were essentially the same with small but overall negligible differences (Figure S8).

### rA_30_ HCG Ensemble Reproduces SAXS Profiles Measured at Different Salt Concentrations After Gentle BioEn Refinement

In their study on the salt dependence of ssRNA rA_30_ and rU_30_, Plumridge et al. ^18^ have shown that the global structure of these highly charged polymers are dependent on (i) the concentration of ions in the solvent and (ii) the ion type. Here, we compared the rA_30_ HCG ensemble, grown from rA_4_ fragments simulated at 150 mM NaCl, to their SAXS profiles measured at 20, 100, 200, 400 and 600 mM NaCl concentration. Despite the fact that we grew the polymer via HCG without taking into account long-range electrostatics beyond the fragment level, the HCG profile matched experimental SAXS profiles recorded at different salt concentrations reasonably well even without reweighting (Figures S9-S13A blue).

BioEn reweighting of the HCG ensemble against the scattering profiles measured at the respective salt concentration established nearly perfect agreement with the experimental profiles (Figure 4A, Figure S9-S13A orange). For each salt concentration we chose a set of weights for the regularisation parameter *θ* = 100 resulting in red. *χ*^2^ *<* 2. According to the L-curve analysis, with Kullback-Leibler divergences close to zero and the cumulative distribution functions (CDF) of rank-ordered weights staying close to uniform reference weights, all BioEn reweightings placed a rather gentle bias on the initial ensemble (Figure S6).

**Figure 4:**
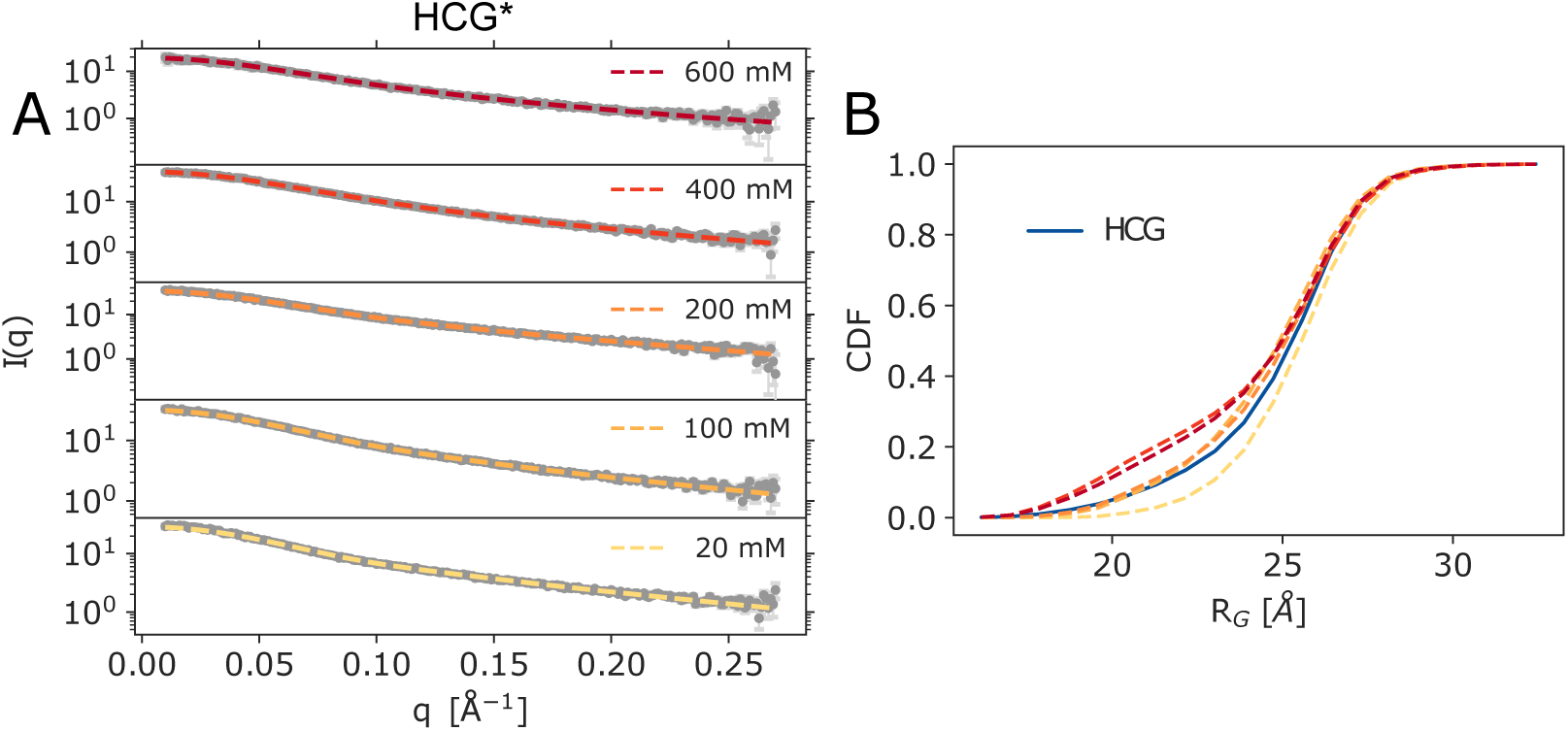
SAXS measurements of rA_30_ at different salt concentrations. (A) SAXS profiles from the rA_30_ HCG ensemble refined against experimental profiles measured at 20, 100, 200, 400, and 600 mM NaCl, HCG* shown in yellow to dark red. Experimental profiles are shown in gray, errors are shown in light gray. (B) Cumulative distribution of *R_G_* as predicted for HCG using Crysol^64^ in blue and the weighted distributions using the refined weights.

The reweighted RMS *R_G_* values for the rA_30_ ensemble were shifted towards the experimental values for each concentration (Figures S9-S13C). The shape of the *R_G_* distributions was minimally modified when we applied the refined weights for 20, 100 and 200 mM (Figures 4B left column and S9, S10, and S11C). In fact, for 100 and 200 mM NaCl, the RNA conformations of rA_30_ with *R_G_ <* 20 Å lost weight against conformations with 20Å *< R_G_ <* 25 Å (Figure 3B left column and S11C). For the highest salt concentrations of 400 and 600 mM NaCl, a distinct shoulder developed in the reweighted *R_G_* distribution at *R_G_ ≈* 20 Å (Figures S12 and S13C). The diminished role of electrostatic repulsion between ssRNA phosphate groups at high salt may explain this trend to compaction.

We assessed the quality of the fit by performing a hplusminus analysis.^65^ The hplusminus test statistic assesses the quality of a fit to ordered data. Applied to the scattering intensities *I*(*q*), it tends to pick up indications for systematic errors, e.g., as a result of deviations in the global size and shape of the ensemble members. Here, going in the HCG* ensembles from low to high salt concentration, we found that systematic deviations at small *q* values decreased, as judged by the residuals and a screening of the signs thereof. In return, we found improved *p*-values for the reweighted ensembles at higher salt concentrations (Figures S9-S13A and B).

Despite the overall efficient reweighting of the HCG ensemble resulting in almost perfect agreement with experiment, we emphasize that the refined ensemble lacks information on electrostatics and other interaction. For a more detailed assessment, one could perform additional MD simulations at the respective salt condition using a small subset of the models sampled in the HCG ensemble as start structure and choosing a reasonable force-field.^28,43^ Such simulations would provide information on electrostatic interactions and the solvent layer.

### HCG Ensembles of rUCAAUC Capture SAXS and NMR Experiments

Using the heterotetramer fragment library for HCG, we are able to grow heteropolymeric ssRNAs of arbitrary sequence. In the following, we show results for the rUCAAUC hexamer, which has been investigated previously by experiments and MD simulations.^14,66^ Visually, the structures appeared rather extended, albeit with populations of structures in which one or two nucleotides were unstacked, similar to what was observed by Bergonzo et al. ^14^ (Figures 2 and 5A).

**Figure 5:**
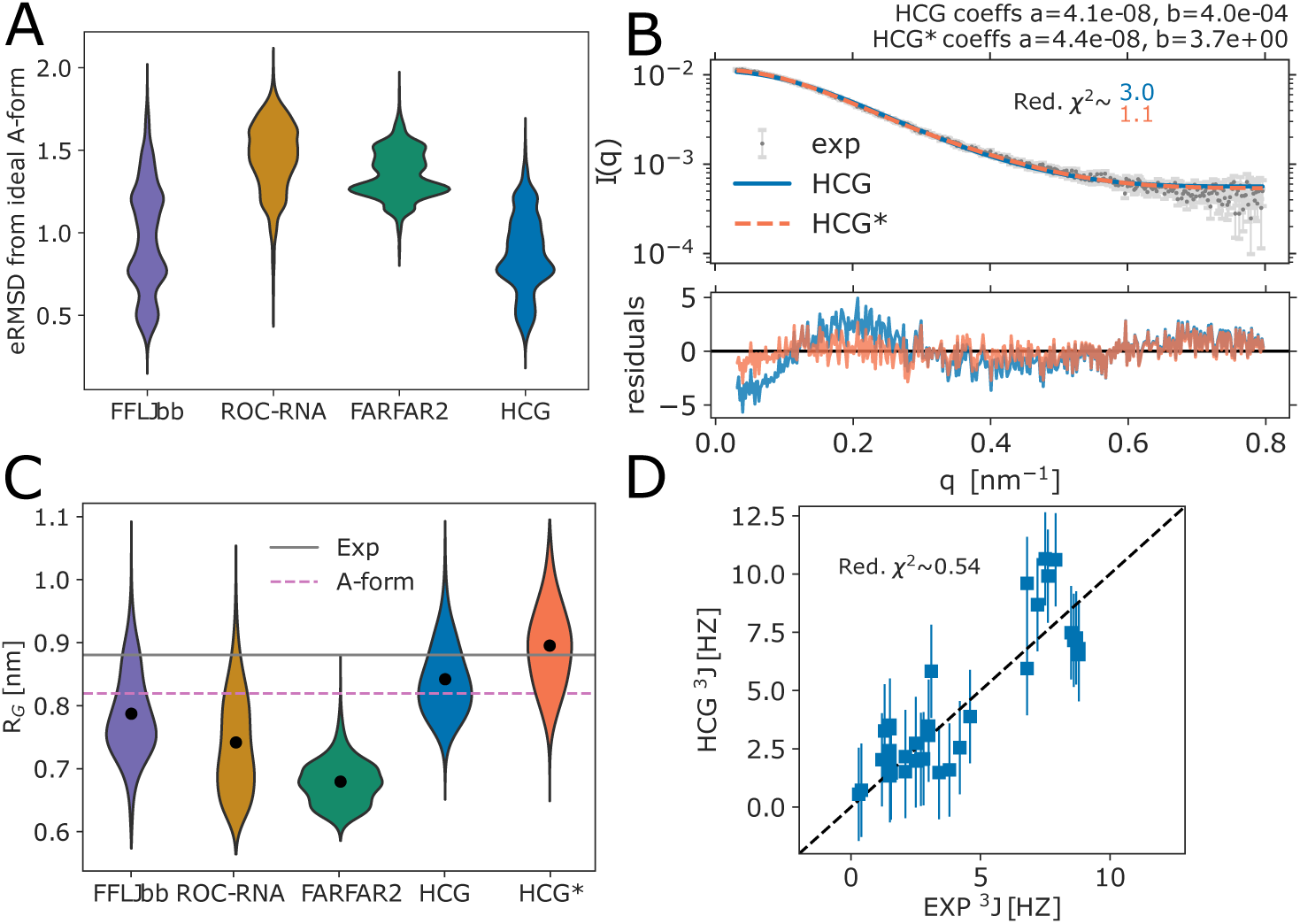
Characterization of structural ensembles of heteropolymeric UCAAUC sampled with MD and FARFAR2 from Ref.,^14^ and HCG. (A) Distribution of the eRMSD to ideal A-form. (B) Top: Average SAXS profile calculated for the HCG ensemble before (blue) and after ensemble refinement (HCG* with weights for *θ* = 100 in orange) fitted to the experimental profile^14^ (orange and gray, respectively). The intensity scale factors a and the background correction constants b as calculated by least-square fitting are shown in the plot. Bottom: Residuals calculated for scattering profiles. (C) Distribution of R*_G_* values in sampled ensembles, the refined HCG* ensemble, experimental average as determined with SAXS, and for typical A-form. (D) Correlation plot of experimentally measured ^3^*J* couplings of the backbone and sugar moiety^66^ and calculated values for HCG. Error estimates for the HCG ensemble of 2 Hz are indicated as lines.

Judging from the comparison to the published SAXS data,^14^ the HCG ensemble of rUCAAUC chains captured the global dimensions without any refinement. The average SAXS profile calculated for the HCG ensemble was in good agreement with the experimental profile, with a reduced *χ*^2^ of about 3.0 and small deviations at small *q* (Figure 5B). For reference, Bergonzo et al.^14^ found profiles of similar quality in their MD simulations of full-length rUCAAUC. For the LJbb force field, their agreement with the SAXS experiments was slightly better, with *χ*^2^ *≈* 2.4 and the deviations at small *q* being less pronounced.

The distribution of *R_G_*in the HCG ensemble was in line with the distribution in the MD ensemble sampled with the LJbb force field (FFLJbb) by Bergonzo et al.^14^ (Figure 5C). The RMS R*_G_* as calculated for the HCG ensemble was slightly closer to the experimentally determined value than the ensemble from full MD simulations. Interestingly, the RMS *R_G_*is close to that of an rUCAAUC polymer in ideal A-form helix conformation. Overall the conformations sampled with HCG seemed to resemble a typical A-form to a larger extent than conformations sampled with the other approaches shown here, with the eRMSD from a typical A-form being smaller on average (Figure 5A).

The analysis of NMR ^3^*J* couplings of the backbone and sugar moiety revealed that overall HCG sampled local properties, as reflected in the torsion angles of the sugar moiety as well as the nucleic acid backbone, in excellent agreement with experiments (Figure 5D). Small deviations in the calculated ^3^*J* couplings were within their predicted uncertainty (*≈*2 HZ for the Karplus relation used to calculate the scalar coupling^55^), resulting in a reduced *χ*^2^ value of *≈*0.54. Thus, we found our ensemble to agree with experiment as good as the MD ensemble (LJbb forcefield) from Bergonzo et al.^14^ We note that for the experimental values, Zhao et al. suggested an error of 2 Hz as well due to the deviations of measured values in multiple independent measurements (see Ref.^66^ Table S4), which we did not consider here.

We reweighted the SAXS profiles calculated for HCG against the experimentally measured scattering profile. Using a small bias with weights for *θ* = 100 we found almost perfect agreement with the experimental profile with reduced *χ*^2^ *≈* 1.1 and *S*_KL_ *«* 1 (Figure S14A, B). Deviations we observed for the refined profile were within the experimental error range and only very small deviations for *q <* 0.1 nm resulted in residuals below 0. The refined weights were used to calculate weighted distributions and averages for properties we analyzed here. The weighted distribution of *R_G_* values was shifted towards larger values with the weighted RMS *R_G_* value being in perfect agreement with the experimental value. Interestingly, we observed only small and overall negligible changes for the eRMSD from ideal A-form and scalar couplings (Figure S14C, D).

### HCG Grows Ensembles with a Large Conformational Variability

Interestingly, we observed a high diversity of global dimensions although we use rather short input fragments (e.g., rU_30_ grown from heteropolymeric tetramers Figure 3, right panel). Using larger fragment sizes to prepare a fragment library, e.g, pentamers with the central trimers and the 3^1^ terminal capping nucleotide being flexible and the 5^1^ terminal cap fixed, would still be computationally feasible, with 4^4^ = 256 fragments. It is interesting to speculate if we may be able to sample a higher population of structures that feature important local motifs, by sampling more local interactions within the input MD fragments.

We compared the sampled structural diversity in HCG and MD ensembles in terms of pairwise RMSDs, calculated using all heavy atoms within a polymer. For short chains (tetramers and hexamers) the distribution of the pairwise RMSD within MD ensembles was slightly larger than within the HCG ensemble. This suggests that a slightly larger conformational variability was sampled with MD (Figure S15), at least in terms of the pairwise RMSD. The pairwise RMSDs between MD and HCG was distributed around 4 nm (Figure S15, rose), similar to what we observed for two independent HCG ensembles of the same polymer (Figure S15C, dashed gray). For the rA_4_ all distributions were shifted towards smaller pairwise RMSDs, probably due to the large population of A-form like helix conformations (Figures S1 and S15A).

Importantly, we do not know the actual extent of structural variability and the expected distribution of pairwise RMSDs for a native ssRNA ensemble. For ensemble refinement, however, it is advantageous to have a broad sampling that covers the relevant conformation space. By integrating experimental information, ensemble refinement methods such as BioEn^37,38^ then down-weight conformations with low statistical relevance. By contrast, if the starting ensemble does not cover the relevant conformation space, conformations in this region would have to be added by biased sampling for a proper ensemble refinement.

In general, efficient comparisons of structurally heterogeneous ensembles is difficult. ^27^ Several algorithms exist to cluster ensemble members according to different properties, often accompanied by machine learning techniques. However, finding appropriate collective variables that really capture the important properties needed to display the differences between ensembles is not straightforward. Recently, a tool to compare structural ensembles of IDPs by determining differences in distributions of local and global properties of the conformations based on a Wasserstein metric was introduced.^70^

### ssRNA Polymers Grown with HCG can be Combined with dsRNA

Conformations sampled with HCG can easily be combined with structured dsRNA or other ordered structures. Here, we exemplarily modeled a region of the 5^1^ UTR of the SARS-CoV-2 genomic RNA (sequence shown in Figure S2). Structured stem-loops were taken from earlier studies using FARFAR2^53,71^ and from NMR studies.^54^ To model the 5^1^ UTR we used MD trajectories of five stem-loops provided by Bottaro et. al^53^ as input ensembles for the structured parts (see Methods). The stem-loop, highlighted in red, were then connected by single-stranded regions grown with HCG, highlighted in blue, resulting in a model containing 233 nucleotides in total. The final ensemble with about 44 different conformations, features models with more extended and more compact single-stranded regions, dictating the overall global dimensions of the modeled 5^1^ UTR (see Figure 6A). For illustration, we randomly chose two structures (see Figure 6 B and C, respectively).

**Figure 6:**
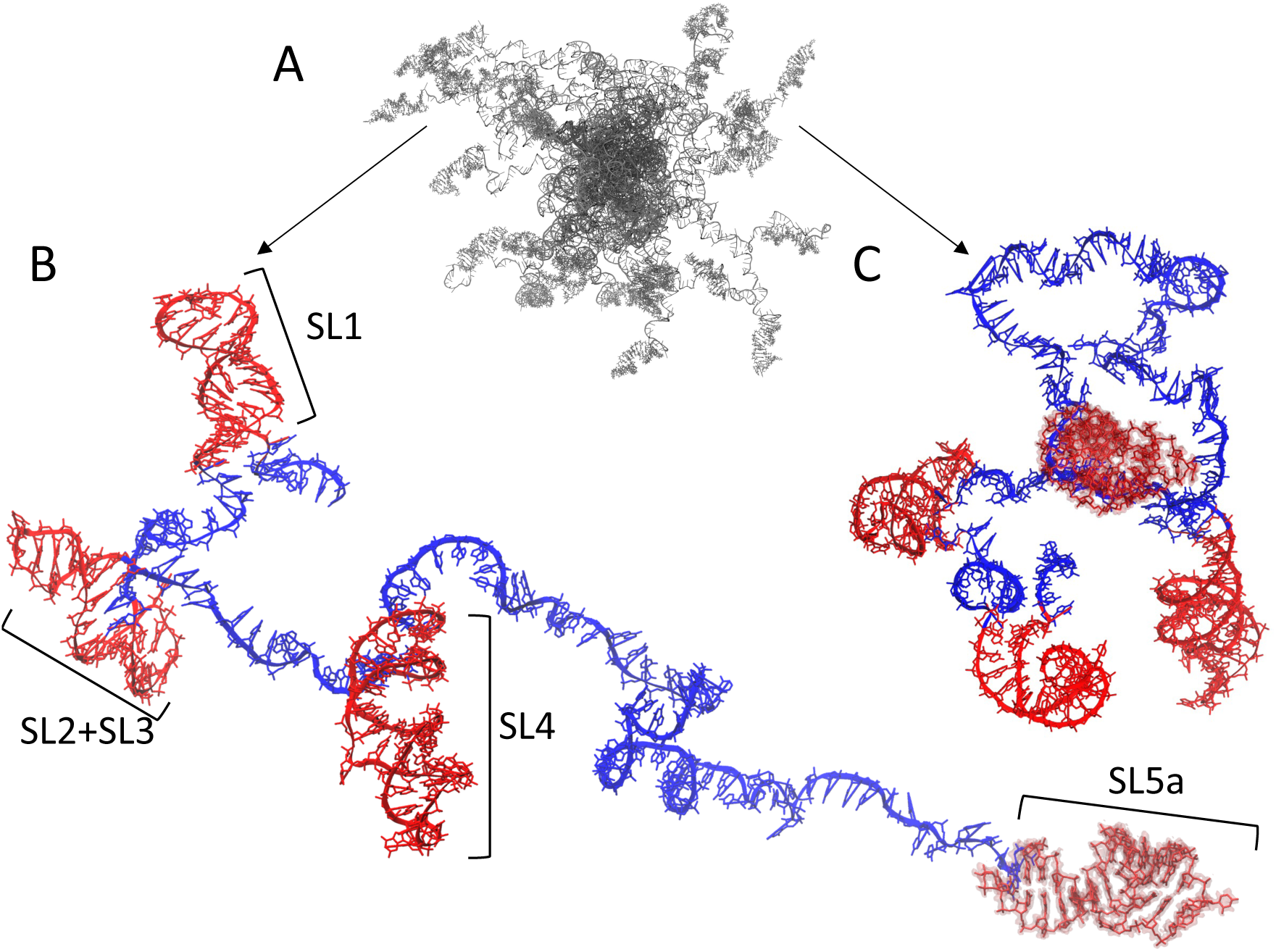
Model of the 5^1^ UTR region of SARS-CoV-2 genomic RNA built by HCG. (A) Ensemble overview of the 5 . (B) and (C) show two representative of extended and compact conformations, respectively, with more detail. Structures of the five stem-loop regions drawn at random from MD trajectories^53^ are shown in red. The connecting single-stranded RNA is shown in blue. The structures are shown with atomic detail. Hydrogen atoms are omitted for clarity. The backbone atoms are shown in a cartoon representation except for the last stem-loop at the 3^1^ end, which is highlighted as a surface. The full-length structure shown here covers 233 nucleotides.

We here demonstrated a possible application of HCG to model structures of RNA molecules that combined structured and unstructured regions such as mRNA molecules. More generally, a similar scheme may be applied to model any kind of biomolecule featuring unstructured parts, e.g., by adding a polyA tail to mRNA. Importantly, since HCG is modular, we can either add additional assembly steps and assemble the different regions after the initial growth or grow the flexible chain with HCG directly at the structured biomolecule. In particular, such models can be used for further analysis, e.g., as initial structure for MD simulations.

### HCG Ensembles of rA_19_ with Mapped Dyes are Somewhat too Extended on Average as Judged by Experimental FRET Efficiencies

We calculated FRET efficiencies for the HCG ensemble of rA_19_ and compared them to the measured mean FRET efficiency *(E)* = 0.56 *±* 0.03 obtained in single-molecule FRET experiments at 150 mM NaCl concentration.^21^ For model 1 with fast and isotropic averaging for the dye orientations about fixed dye positions (eq. 1), we obtained a mean efficiency of *(E) ≈* 0.41. In model 2 we start from the ensemble in model 1, but with multiple dye pair conformations placed onto every conformation *i* in the rA_19_ ensemble. By averaging the FRET efficiency over these dye pairs and their orientations, we effectively assumed dynamic dyes in model 2, which we consider to be more realistic than model 1. We observed a mean FRET efficiency of *(E) ≈* 0.42. In the less realistic model 3 with fully static dyes, we fixed inter-dye distances and determined *κ*^2^ explicitly from the dye conformations (eq. 3). For model 3, we obtained *(E) ≈* 0.32 with a high population of conformations having *E <* 0.1. By comparison, MD simulations of full-length rA_19_ using the same force field as in our fragment MD simulation gave FRET efficiencies of *(E) ≈* 0.3,^21^ calculated with explicit dyes mapped onto the sampled conformers and *κ*^2^ = 2*/*3 fixed, as in our model 1. Differences we observed for FRET efficiencies calculated from the MD and the HCG ensemble using model 1 may indicate that the MD simulation was too short with 7 *µ*s of sampling in aggregate. Alternatively, we may have a favorable compensation of errors in chain growth by accounting primarily for the local structure.

Using BioEn,^37,38^ we then gently reweighted the ensembles of models 1, 2, and 3 to match the experimental mean FRET efficiency. For model 1 and model 2, we obtained reduced *χ*^2^ *≈* 1.4 for a BioEn confidence parameter *θ* = 40, and for model 3 we obtained *χ*^2^ *≈* 2.3 for *θ* = 60 (see Figure 7A orange top row, middle row, and bottom row, respectively). The ensemble refinement assigned higher weights to the tail of the ensemble, i.e., to more compact chains with *E »* 0.6. In turn, the weighted distributions and the average inter-dye distance were shifted to shorter distances and with that within the range of the inter-dye distance as inferred from the single-molecule FRET experiment using a worm-like chain polymer model (see Figure 7B). Here, either the structures were more compact, the mapped dyes featured less extended linkers and/or the mapped dyes pointed towards each other (Figure S3). This shift in population toward more compact structures is qualitatively consistent with what we found in the BioEn reweighting for rA_30_ according to the SAXS data (Figure 3B. However, the shift there was considerably smaller, as the *R_G_* value before already agreed with the measurements within the uncertainty. We note that the *r* distributions in the HCG* ensembles for models 1 and 3 are nearly identical (Figure 7B). For model 2, the distance distribution in HCG was narrower and for HCG* the peak of the distribution was slightly shifted towards larger inter-dye distances. A small shoulder at around 4 nm dye-dye distance, present in HCG* for all models, was more pronounced.

**Figure 7:**
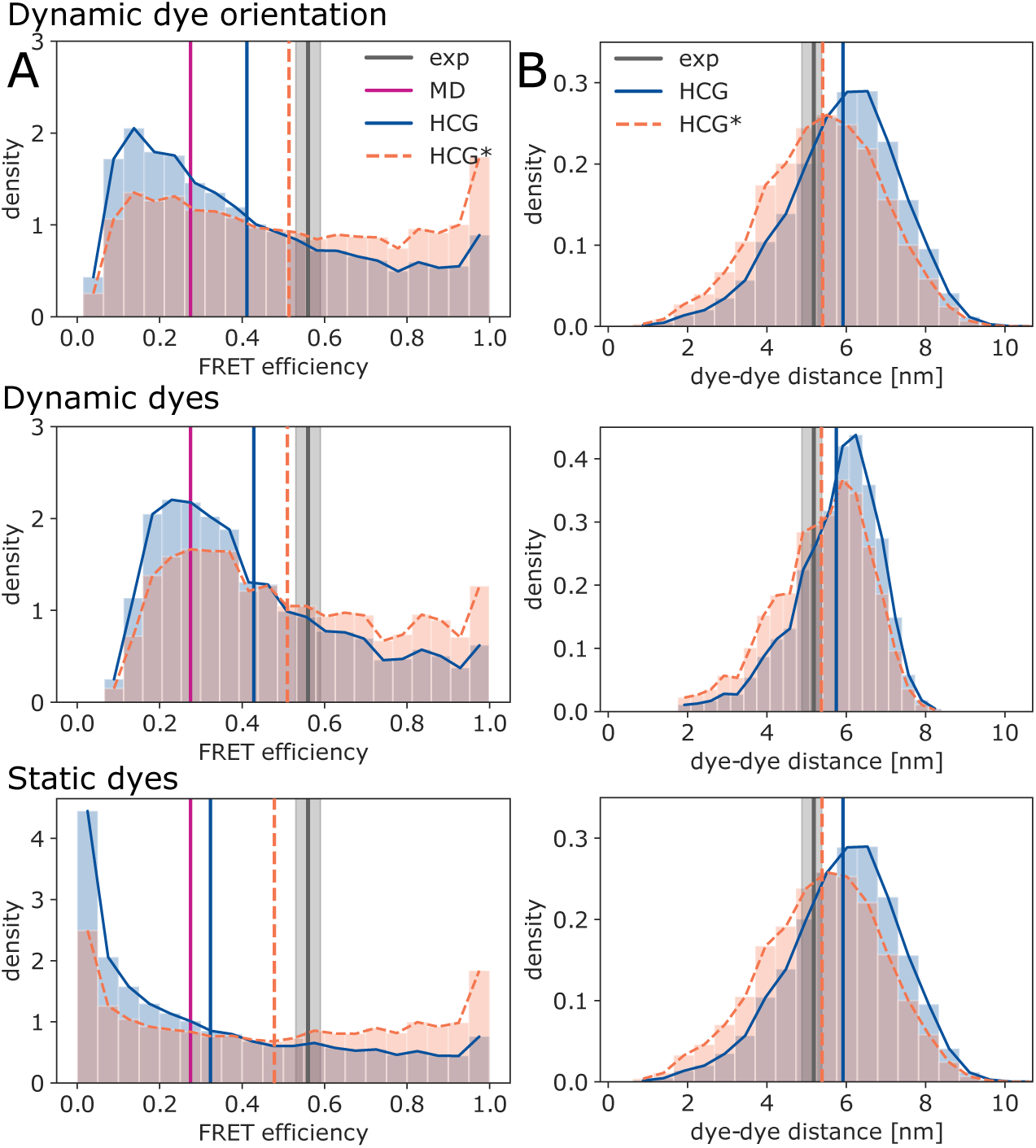
HCG ensembles of rA_19_ compared to single-molecule FRET experiments. (A) Distribution of FRET efficiencies in the HCG ensemble (blue) and in the reweighted HCG* ensemble (orange) calculated with model 1 (top) with dynamic dye orientations and *κ*^2^ = 2*/*3, model 2 (middle) with dynamic dyes and *κ*^2^ = 2*/*3, and model 3 (bottom) with static dyes. For HCG* we chose refined weights for *θ* = 40 (models 1 and 2) and *θ* = 60 (model 3). Vertical lines indicate the mean FRET efficiency measured in experiment (black, with gray shading indicating *±*SEM), sampled in MD simulations of full-length rA_19_ with the same force field as used here to build fragment libraries^21^ (magenta), and calculated for the HCG (blue) and HCG* ensembles (orange). (B) Distributions of the inter-dye distance as determined for the HCG ensemble (blue) and the reweighted HCG* ensemble (orange) with models 1 (top), 2 (middle), and 3 (bottom). The mean distances for experiment (black), MD simulation (magenta), HCG (blue), and HCG* (orange) are shown as vertical solid and dotted lines. The experimental mean inter-dye distance was inferred from experimental single-molecule FRET efficiencies using a worm-like-chain model for the distance probability density function.^21^

The shape of the reweighted distributions of the FRET efficiency may indicate slight overfitting. However, judging from the L-curve analysis and the CDF of rank-ordered weights (Figure S16, orange, dark green, and dark red), the set of weights we chose seemed to impose a rather gentle bias with *S*_KL_ *<* 0.2 for all three models. An important point to consider is how to properly perform the ensemble reweighting for polymers with attached labels.^39,72^ In approaches such as FRETpredict^73^ dyes are placed onto proteins using a rotamer library approach (RLA) in order to predict FRET efficiencies. Here, the dyes are given individual statistical weights. In the present study we reweighted the whole molecule, i.e., polymer chain plus the attached fluorophore molecules. As an alternative, one could reweight the chain and dye separately.

## 4 CONCLUSIONS

Single-stranded RNA appears prominently in many cellular regulation processes, e.g., in mRNA and its poly-A tail but also in loops and linkers. The structural modeling of flexible nucleic acid with unpaired nucleobases poses formidable challenges. Here, we showed that hierarchical chain growth, previously introduced for disordered proteins,^43^ can be used to produce structural ensembles of ssRNA with atomic detail, starting from fragments sampled in MD simulations. The resulting structural ensembles feature highly diverse conformations (Figures 2 and S3) in good agreement with NMR experiments probing the local structure (Figure 5D). Also SAXS and FRET experiments probing the global structure are reproduced well. Overall, we found the HCG ensembles to agree with experiment about as well or better than MD simulations of full-length ssRNA (Figure 3-5, 7).

HCG relies on a number of simplifying assumptions. Most importantly, it assumes that the relevant local structure of disordered biopolymers is sampled properly in short fragments and that these fragments can be assembled into full length chains subject to only steric interactions. In particular, in its simplest form HCG does not account for long-range electrostatic interactions. It is therefore remarkable that we obtained excellent agreement for rA_30_ SAXS data over a wide range of salt concentrations, in particular for 100-200 mM NaCl (Figure 4). The computational efficiency of HCG makes it possible to construct large ensembles with diverse conformations, sampling also significant populations of rare but relevant conformations. This broad coverage of conformation space enables ensemble reweighting schemes to match a wide range of experiments within expected uncertainties (Figure 3-5, 7, S6, S7, S9-S13, S14A and B, and S16).

HCG is implemented as a Monte Carlo chain growth algorithm with a well-defined ensemble and partition function.^43^ Therefore, HCG can easily be combined with other Monte Carlo sampling techniques, e.g., to perform importance sampling as in the reweighted hierarchical chain growth (RHCG).^40^ In RHCG, one uses a fragment library that is refined against experimental data prior to fragment assembly to improve the sampling of local properties in the grown full-length ensemble. An exciting perspective is to adopt an RHCG-like sampling scheme to include information on inter-fragment interactions during chain growth or to grow loop structures. This task may be turned into a machine learning problem. Methods based on artificial intelligence (AI) have been proven to reliably predict tertiary structure of folded double-stranded and also single-stranded RNA.^30,71,74,75^ Query sequences that require modeling of both structured and disordered regions may be an excellent target for AI-guided applications of Monte Carlo techniques. We have shown that HCG is suited to model segments of mRNA that feature structured and unstructured regions (see Figure 6). HCG in combination with machine learning approaches could prove useful for modeling more complicated mRNA or long-non-coding RNA (lncRNA) with internal short disordered loops. To improve the grown structures one can include experimentally derived information^40^ and information from secondary structure prediction tools. Fragment libraries for 3D structures of RNA secondary structure motifs^76^ can be used as input for HCG of more complex RNA folds. One could also use coarse-grained RNA simulation models^77^ to build fragment libraries for the assembly of large RNA structures.

HCG can also be combined with MD simulations to gain insight on inter- and intramolecular interactions and the dynamics of ssRNA. In previous work, we have shown that the conformations of IDPs sampled with HCG are perfectly suited as starting structures for parallel but independent MD simulations with atomic detail.^43^ Similarly, this could be done with the ssRNA conformations as modeled here, either the fully flexible single chains or molecules with structured and flexible regions. Starting from a multitude of reasonable initial structures will facilitate the exploration of the relevant conformational space and dynamics.

## Supporting Information Available

Supporting figures of: Cluster analysis of the MD trajectory of a rA_4_ fragment, sequence of the SARS-CoV-2 5^1^ UTR as modelled here; snapshots of rA_19_ models from HCG with attached FRET labels; stacking analysis of poly A and poly U 30mers from HCG; backbone dihedral angle distributions sampled in rA_4_ MD trajectories and rA_19_ HCG models; BioEn reweighting of rA_30_ HCG ensemble against SAXS data at different salt concentrations; BioEn reweighting of rU_30_ HCG ensemble against experimental SAXS data; distribution of the radius of gyration for HCG rA_30_ ensembles from homo- or heteropolymers; BioEn reweighting of rUCAAUC HCG ensemble against experimental SAXS data; CDF of pairwise RMSDs in (RE)MD ensembles and HCG ensembles of ssRNA polymers with different lengths; BioEn reweighting of rA_19_ HCG ensemble against experimental FRET data.

## ACKNOWLEDGEMENTS

We thank Kara Grotz for sharing her data and insightful discussions. We thank Mark Nüesch and Benjamin Schuler for comments and detailed exchange on the analysis of the singlemolecule FRET data. We thank Jürgen Köfinger for fruitful discussion and for sharing his experience with the analysis of scattering data, ensemble refinement via Bayesian Inference of Ensembles, and the hplusminus analysis.

## Graphical TOC Entry

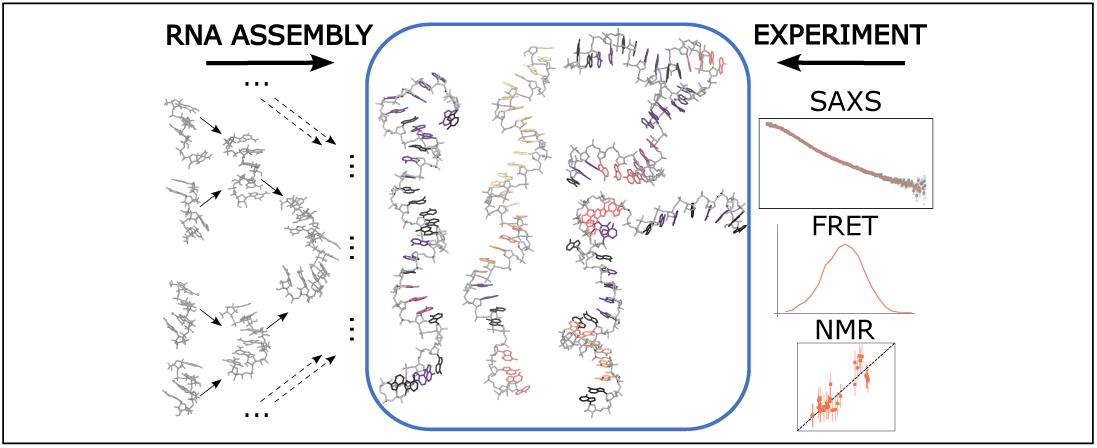

## References

(1) Scull, C. E.; Dandpat, S. S.; Romero, R. A.; Walter, N. G. Transcriptional riboswitches integrate timescales for bacterial gene expression control. Front. Mol. Biosci. 2021, 7 .

(2) Li, Q. Q.; Liu, Z.; Lu, W.; Liu, M. Interplay between alternative splicing and alternative polyadenylation defines the expression outcome of the plant unique OXIDATIVE TOLERANT-6 Gene. Sci. Rep. 2017, 7 .

(3) Passmore, L. A.; Coller, J. Roles of mRNA poly(A) tails in regulation of eukaryotic gene expression. Nat. Rev. Mol. Cell Biol. 2022, 23, 93–106.

(4) Leulliot, N.; Varani, G. Current topics in RNA-protein recognition: control of pecificity and biological function through induced fit and conformational capture. Biochemistry 2001, 40, 7947–7956.

(5) Ku, T. H.; Zhang, T.; Luo, H.; Yen, T. M.; Chen, P. W.; Han, Y.; Lo, Y. H. Nucleic acid aptamers: An emerging tool for biotechnology and biomedical sensing. Sens. Switz. 2015, 15, 16281–16313.

(6) Eulalio, A.; Huntzinger, E.; Izaurralde, E. Getting to the root of miRNA-mediated gene silencing. Cell 2008, 132, 9–14.

(7) Sahin, U.; Karikό, K.; Türeci, Ö. MRNA-based therapeutics-developing a new class of drugs. Nat. Rev. Drug Discov. 2014, 13, 759–780.

(8) Damase, T. R.; Sukhovershin, R.; Boada, C.; Taraballi, F.; Pettigrew, R. I.; Cooke, J. P. The limitless future of RNA therapeutics. Front. Bioeng. Biotechnol. 2021, 9 .

(9) Khurana, A.; Allawadhi, P.; Khurana, I.; Allwadhi, S.; Weiskirchen, R.; Banothu, A. K.; Chhabra, D.; Joshi, K.; Bharani, K. K. Role of nanotechnology behind the success of mRNA vaccines for COVID-19. Nano Today 2021, 38 .

(10) Eichhorn, C. D.; Feng, J.; Suddala, K. C.; Walter, N. G.; Brooks, C. L.; AlHashimi, H. M. Unraveling the structural complexity in a single-stranded RNA tail: Implications for efficient ligand binding in the prequeuosine riboswitch. Nucleic Acids Res. 2012, 40, 1345–1355.

(11) Tubbs, J. D.; Condon, D. E.; Kennedy, S. D.; Hauser, M.; Bevilacqua, P. C.; Turner, D. H. The nuclear magnetic resonance of CCCC RNA reveals a right-handed helix, and revised parameters for AMBER force field torsions improve structural predictions from molecular dynamics. Biochemistry 2013, 52, 996–1010.

(12) Condon, D. E.; Kennedy, S. D.; Mort, B. C.; Kierzek, R.; Yildirim, I.; Turner, D. H. Stacking in RNA: NMR of four tetramers benchmark molecular dynamics. J. Chem. Theory Comput. 2015, 11, 2729–2742.

(13) Bottaro, S.; Bussi, G.; Kennedy, S. D.; Turner, D. H.; Lindorff-Larsen, K. Conformational ensembles of RNA oligonucleotides from integrating NMR and molecular simulations. Sci. Adv. 2018, 4 .

(14) Bergonzo, C.; Grishaev, A.; Bottaro, S. Conformational heterogeneity of UCAAUC RNA oligonucleotide from molecular dynamics simulations, SAXS, and NMR experiments. RNA 2022, 28, 937–946.

(15) Liu, B.; Shi, H.; Al-Hashimi, H. M. Developments in solution-state NMR yield broader and deeper views of the dynamic ensembles of nucleic acids. Curr. Opin. Struct. Biol. 2021, 70, 16–25.

(16) Chen, H.; Meisburger, S. P.; Pabit, S. A.; Sutton, J. L.; Webb, W. W.; Pollack, L. Ionic strength-dependent persistence lengths of single-stranded RNA and DNA. Proc. Natl. Acad. Sci. 2012, 109, 799–804.

(17) Plumridge, A.; Meisburger, S. P.; Andresen, K.; Pollack, L. The impact of base stacking on the conformations and electrostatics of single-stranded DNA. Nucleic Acids Res. 2017, 45, 3932–3943.

(18) Plumridge, A.; Andresen, K.; Pollack, L. Visualizing disordered single-stranded RNA: connecting sequence, structure, and electrostatics. J. Am. Chem. Soc. 2020, 142, 109– 119.

(19) Schuler, B.; Eaton, W. A. Protein folding studied by single-molecule FRET. Curr. Opin. Struct. Biol. 2008, 18, 16–26.

(20) Zheng, W.; Borgia, A.; Buholzer, K.; Grishaev, A.; Schuler, B.; Best, R. B. Probing the action of chemical denaturant on an intrinsically disordered protein by simulation and experiment. J. Am. Chem. Soc. 2016, 138, 11702–11713.

(21) Grotz, K. K.; Nuuesch, M. F.; Holmstrom, E. D.; Heinz, M.; Stelzl, L. S.; Schuler, B.; Hummer, G. Dispersion correction alleviates dye stacking of single-stranded DNA and RNA in simulations of single-molecule fluorescence experiments. J. Phys. Chem. B 2018, 122, 11626–11639.

(22) Kührová, P.; Mlýnský, V.; Zgarbová, M.; Krepl, M.; Bussi, G.; Best, R. B.; Otyepka, M.; Šponer, J.; Banáš, P. Improving the performance of the amber RNA force field by tuning the hydrogen-bonding interactions. J. Chem. Theory Comput. 2019, 15, 3288–3305.

(23) Tan, D.; Piana, S.; Dirks, R. M.; Shaw, D. E. RNA force field with accuracy comparable to state-of-the-art protein force fields. Proc. Natl. Acad. Sci. U. S. A. 2018, 115, E1346– E1355.

(24) Grotz, K. K.; Cruz-Leόn, S.; Schwierz, N. Optimized magnesium force field parameters for biomolecular simulations with accurate solvation, ion-binding, and water-exchange properties. J. Chem. Theory Comput. 2021, 17, 2530–2540.

(25) Cruz-Leόn, S.; Vanderlinden, W.; Müller, P.; Forster, T.; Staudt, G.; Lin, Y.-Y.; Lipfert, J.; Schwierz, N. Twisting DNA by salt. Nucleic Acids Res. 2022, 50, 5726–5738.

(26) Best, R. B.; Hummer, G. Optimized molecular dynamics force fields applied to the helix-coil transition of polypeptides. J. Phys. Chem. B 2009, 113, 9004–9015.

(27) Ahmed, M. C.; Skaanning, L. K.; Jussupow, A.; Newcombe, E. A.; Kragelund, B. B.; Camilloni, C.; Langkilde, A. E.; Lindorff-Larsen, K. Refinement of *α*-synuclein ensembles sgainst SAXS data: comparison of force fields and methods. Front. Mol. Biosci. 2021, 8 .

(28) Pietrek, L. M.; Stelzl, L. S.; Hummer, G. Structural ensembles of disordered proteins from hierarchical chain growth and simulation. Curr. Opin. Struct. Biol. 2023, 78, 102501.

(29) Das, R.; Karanicolas, J.; Baker, D. Atomic accuracy in predicting and designing noncanonical RNA structure. Nat. Methods 2010, 7, 291–294.

(30) Watkins, A. M.; Rangan, R.; Das, R. FARFAR2: Improved de novo rosetta prediction of complex global RNA folds. Structure 2020, 28, 963–976.e6.

(31) Chojnowski, G.; Zaborowski, R.; Magnus, M.; Bujnicki, J. M. RNA fragment assembly with experimental restraints. bioRxiv 2021, 2021–02.

(32) Rόżycki, B.; Kim, Y. C.; Hummer, G. SAXS ensemble refinement of ESCRT-III CHMP3 conformational transitions. Structure 2011, 19, 109–116.

(33) Boomsma, W.; Tian, P. F.; Frellsen, J.; Ferkinghoff-Borg, J.; Hamelryck, T.; Lindorff-Larsen, K.; Vendruscolo, M. Equilibrium simulations of proteins using molecular fragment replacement and NMR chemical shifts. Proc. Natl. Acad. Sci. U.S.A. 2014, 111, 13852–13857.

(34) Larsen, A. H.; Wang, Y.; Bottaro, S.; Grudinin, S.; Arleth, L.; Lindorff-Larsen, K. Combining molecular dynamics simulations with small-angle X-ray and neutron scattering data to study multi-domain proteins in solution. PLOS Comput. Biol. 2020, 16, e1007870.

(35) Bottaro, S.; Bengtsen, T.; Lindorff-Larsen, K. In Struct. Bioinforma. Methods protoc.; Gáspári, Z., Ed.; Springer US: New York, NY, 2020; pp 219–240.

(36) Borgia, A.; Zheng, W.; Buholzer, K.; Borgia, M. B.; Schüler, A.; Hofmann, H.; Soranno, A.; Nettels, D.; Gast, K.; Grishaev, A., et al. Consistent view of polypeptide chain expansion in chemical denaturants from multiple experimental methods. J. Am. Chem. Soc. 2016, 138, 11714–11726.

(37) Hummer, G.; Köfinger, J. Bayesian ensemble refinement by replica simulations and reweighting. J. Chem. Phys. 2015, 143 .

(38) Köfinger, J.; Stelzl, L. S.; Reuter, K.; Allande, C.; Reichel, K.; Hummer, G. Efficient ensemble refinement by reweighting. J. Chem. Theory Comput. 2019, 15, 3390–3401.

(39) Reichel, K.; Stelzl, L. S.; Koefinger, J.; Hummer, G. Precision DEER distances from spin-label ensemble refinement. J. Phys. Chem. Lett. 2018,

(40) Stelzl, L. S.; Pietrek, L. M.; Holla, A.; Oroz, J.; Sikora, M.; Köfinger, J.; Schuler, B.; Zweckstetter, M.; Hummer, G. Global structure of the intrinsically disordered protein tau emerges from its local structure. JACS Au 2022, 2, 673–686.

(41) Plumridge, A.; Meisburger, S. P.; Pollack, L. Visualizing single-stranded nucleic acids in solution. Nucleic Acids Res. 2017, 45, e66.

(42) Shi, H.; Rangadurai, A.; Assi, H. A.; Roy, R.; Case, D. A.; Herschlag, D.; Yesselman, J. D.; Al-Hashimi, H. M. Rapid and accurate determination of atomistic RNA dynamic ensemble models using NMR and structure prediction. Nat. Commun. 2020, 11, 5531.

(43) Pietrek, L. M.; Stelzl, L. S.; Hummer, G. Hierarchical ensembles of intrinsically disordered proteins at atomic resolution in molecular dynamics simulations. J. Chem. Theory Comput. 2020, 16, 725–737.

(44) Salomon-Ferrer, R.; Case, D. A.; Walker, R. C. An overview of the Amber biomolecular simulation package. Wiley Interdiscip. Rev. Comput. Mol. Sci. 2013, 3, 198–210.

(45) Piana, S.; Donchev, A. G.; Robustelli, P.; Shaw, D. E. Water dispersion interactions strongly influence simulated structural properties of disordered protein states. J. Phys. Chem. B 2015, 119, 5113–5123.

(46) Joung, I. S.; Cheatham, T. E. Determination of alkali and halide monovalent ion parameters for use in explicitly solvated biomolecular simulations. J. Phys. Chem. B 2008, 112, 9020–9041.

(47) Abraham, M. J.; Murtola, T.; Schulz, R.; Páll, S.; Smith, J. C.; Hess, B.; Lindahl, E. GROMACS: High performance molecular simulations through multi-level parallelism from laptops to supercomputers. SoftwareX 2015, 1–2, 19–25.

(48) Hess, B. P-LINCS: A parallel linear constraint solver for molecular simulation. J. Chem. Theory Comput. 2008, 4, 116–122.

(49) Parrinello, M.; Rahman, A. Polymorphic transitions in single crystals: A new molecular dynamics method. J. Appl. Phys. 1981, 52, 7182–7190.

(50) Darden, T.; York, D.; Pedersen, L. Particle mesh Ewald: An N·log(N) method for Ewald sums in large systems. J. Chem. Phys. 1993, 98, 10089–10092.

(51) Patriksson, A.; van der Spoel, D. A temperature predictor for parallel tempering simulations. Phys. Chem. Chem. Phys. 2008, 10, 2073–2077.

(52) Cordero, B.; Gόmez, V.; Platero-Prats, A. E.; Revés, M.; Echeverría, J.; Cremades, E.; Barragán, F.; Alvarez, S. Covalent radii revisited. Dalton Trans. 2008, 2832.

(53) Bottaro, S.; Bussi, G.; Lindorff-Larsen, K. Conformational ensembles of noncoding elements in the SARS-CoV-2 genome from molecular dynamics simulations. J. Am. Chem. Soc. 2021, 143, 8333–8343.

(54) Wacker, A.; Weigand, J. E.; Akabayov, S. R.; Altincekic, N.; Bains, J. K.; Banijamali, E.; Binas, O.; Castillo-Martinez, J.; Cetiner, E.; Ceylan, B., et al. Secondary structure determination of conserved SARS-CoV-2 RNA elements by NMR spectroscopy. Nucleic Acids Res. 2020, 48, 12415–12435.

(55) Bottaro, S.; Bussi, G.; Pinamonti, G.; Reiber, S.; Boomsma, W.; Lindorff-Larsen, K. Barnaba: software for analysis of nucleic acid structures and trajectories. RNA 2019, 25, 219–231.

(56) McGibbon, R. T.; Beauchamp, K. A.; Harrigan, M. P.; Klein, C.; Swails, J. M.; Hernández, C. X.; Schwantes, C. R.; Wang, L.-P.; Lane, T. J.; Pande, V. S. MDTraj: a modern open library for the analysis of molecular dynamics trajectories. Biophys. J. 2015, 109, 1528–1532.

(57) Michaud-Agrawal, N.; Denning, E. J.; Woolf, T. B.; Beckstein, O. MDAnalysis: A toolkit for the analysis of molecular dynamics simulations. J. Comput. Chem. 2011, 32, 2319–2327.

(58) Gowers, R.; Linke, M.; Barnoud, J.; Reddy, T.; Melo, M.; Seyler, S.; Domański, J.; Dotson, D.; Buchoux, S.; Kenney, I., et al. MDAnalysis: A python package for the rapid analysis of molecular dynamics simulations. Proc. 15th Python Sci. Conf. 2016, 98–105.

(59) Schuler, B.; Lipman, E. A.; Steinbach, P. J.; Kumke, M.; Eaton, W. A. Polyproline and the “spectroscopic ruler” revisited with single-molecule fluorescence. Proceedings of the National Academy of Sciences 2005, 102, 2754–2759.

(60) Hellenkamp, B.; Schmid, S.; Doroshenko, O.; Opanasyuk, O.; Kühnemuth, R.; Rezaei Adariani, S.; Ambrose, B.; Aznauryan, M.; Barth, A.; Birkedal, V., et al. Precision and accuracy of single-molecule FRET measurements—a multi-laboratory benchmark study. Nat. Methods 2018, 15, 669–676.

(61) Best, R. B.; Merchant, K. A.; Gopich, I. V.; Schuler, B.; Bax, A.; Eaton, W. A. Effect of flexibility and cis residues in single-molecule FRET studies of polyproline. Proc. Natl. Acad. Sci. U. S. A. 2007, 104, 18964–18969.

(62) Holmstrom, E. D.; Holla, A.; Zheng, W.; Nettels, D.; Best, R. B.; Schuler, B. Accurate transfer efficiencies, distance distributions, and ensembles of unfolded and intrinsically disordered proteins from single-molecule FRET. Methods Enzymol. 2018, 611, 287–325.

(63) Hummer, G.; Szabo, A. Dynamics of the orientational factor in fluorescence resonance energy transfer. J. Phys. Chem. B 2017, 121, 3331–3339.

(64) Svergun, D.; Barberato, C.; Koch, M. H. J. CRYSOL – a program to evaluate X-ray solution scattering of biological macromolecules from atomic coordinates. J. Appl. Crystallogr. 1995, 28, 768–773.

(65) Köfinger, J.; Hummer, G.; Köfinger, J. Powerful statistical tests for ordered data. ChemRxiv 2021,

(66) Zhao, J.; Kennedy, S. D.; Berger, K. D.; Turner, D. H. Nuclear magnetic resonance of single-stranded RNAs and DNAs of CAAU and UCAAUC as benchmarks for molecular dynamics simulations. J. Chem. Theory Comput. 2020, 16, 1968–1984.

(67) Sripakdeevong, P.; Kladwang, W.; Das, R. An enumerative stepwise ansatz enables atomic-accuracy RNA loop modeling. Proc. Natl. Acad. Sci. 2011, 108, 20573–20578.

(68) Cruz-Leόn, S.; Grotz, K. K.; Schwierz, N. Extended magnesium and calcium force field parameters for accurate ion–nucleic acid interactions in biomolecular simulations. J. Chem. Phys. 2021, 154, 171102.

(69) Chen, Y.; Pollack, L. SAXS studies of RNA: structures, dynamics, and interactions with partners. Wiley Interdiscip. Rev. RNA 2016, 7, 512–526.

(70) González-Delgado, J.; Sagar, A.; Zanon, C.; Lindorff-Larsen, K.; Bernadό, P.; Neuvial, P.; Cortés, J. WASCO: A Wasserstein-based statistical tool to compare conformational ensembles of intrinsically disordered proteins. J. Mol. Biol. 2023, 168053.

(71) Zhang, K.; Zheludev, I. N.; Hagey, R. J.; Haslecker, R.; Hou, Y. J.; Kretsch, R.; Pin-tilie, G. D.; Rangan, R.; Kladwang, W.; Li, S., et al. Cryo-EM and antisense targeting of the 28-kDa frameshift stimulation element from the SARS-CoV-2 RNA genome. Nat. Struct. Mol. Biol. 2021, 28, 747–754.

(72) Tesei, G.; Martins, J. M.; Kunze, M. B. A.; Wang, Y.; Crehuet, R.; Lindorff-Larsen, K. DEER-PREdict: Software for efficient calculation of spin-labeling EPR and NMR data from conformational ensembles. PLOS Comput. Biol. 2021, 17, e1008551.

(73) Montepietra, D.; Tesei, G.; Martins, J. M.; Kunze, M. B. A.; Best, R. B.; Lindorff-Larsen, K. FRETpredict: A Python package for FRET efficiency predictions using rotamer libraries. bioRxiv 2023,

(74) Townshend, R. J. L.; Eismann, S.; Watkins, A. M.; Rangan, R.; Karelina, M.; Das, R.; Dror, R. O. Geometric deep learning of RNA structure. Sci. 80- 2021, 373, 1047–1051.

(75) Baek, M.; McHugh, R.; Anishchenko, I.; Baker, D.; DiMaio, F. Accurate prediction of nucleic acid and protein-nucleic acid complexes using RoseTTAFoldNA. bioRxiv 2022,

(76) Gan, H. H.; Pasquali, S.; Schlick, T. Exploring the repertoire of RNA secondary motifs using graph theory; implications for RNA design. Nucleic Acids Res. 2003, 31, 2926– 2943.

(77) Cragnolini, T.; Laurin, Y.; Derreumaux, P.; Pasquali, S. Coarse-Grained HiRE-RNA Model for ab Initio RNA Folding beyond Simple Molecules, Including Noncanonical and Multiple Base Pairings. J. Chem. Theory Comput. 2015, 11, 3510–3522.

