## Supplementary figures S1-S16 for "Hierarchical Assembly of Single-Stranded RNA"

### Supporting Figures

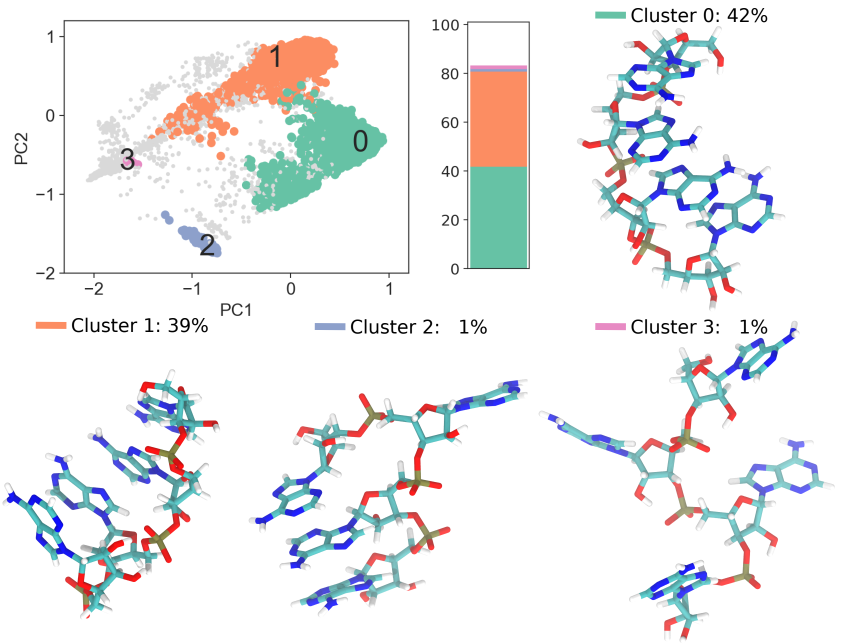


Figure S1: Cluster analysis of the MD trajectory of a $rA_4$ fragment. A) PCA of the g-vectors of 10000 conformtions sampled in 100 ns REMD. The conformations are assigned to six different clusters. The four most populated clusters are highlighted in color and a representative structure for each cluster is shown as snapshot in (B-E). The cluster analysis was performed using the Barnaba package.[S1]

SL1

1 AUUAAGGGUU UAUACCUUCC CAGGUAACAA ACCCAACCAA CUUUCGGAUC

SL1+SL3

51 UCUUGUAGAU CUGUUCUCUA AACGAACGAA CUUUAAAAUC UGGGUGUGGC

SL4

101 UGUCACUCGG CUGCAUGCUU AGUGCACUCA CGCCCAGUAU AAUUAAUAAC

151 UAAUUUUACU GUCGUUGACA GGACACGAGU AACUCGUCUA UCUUCUGCAG

SL5a

201 GGCUGCUUAC GGUUUCGUCC GUGUUGCAGC CCC

Figure S2: Sequence of the SARS-CoV-2 5′ UTR as modelled here. Base paired stem-loop (SL) regions as sampled with MD, published previously,[S2] are highlighted in red, single-stranded regions modelled with HCG, are highlighted in blue. Nucleobases 162-200 are part of SL5, i.e., are predicted to be structured but in our example this region is unstructured, modelled with HCG (underlined in red). Sequence information was taken from Ref.[S2]

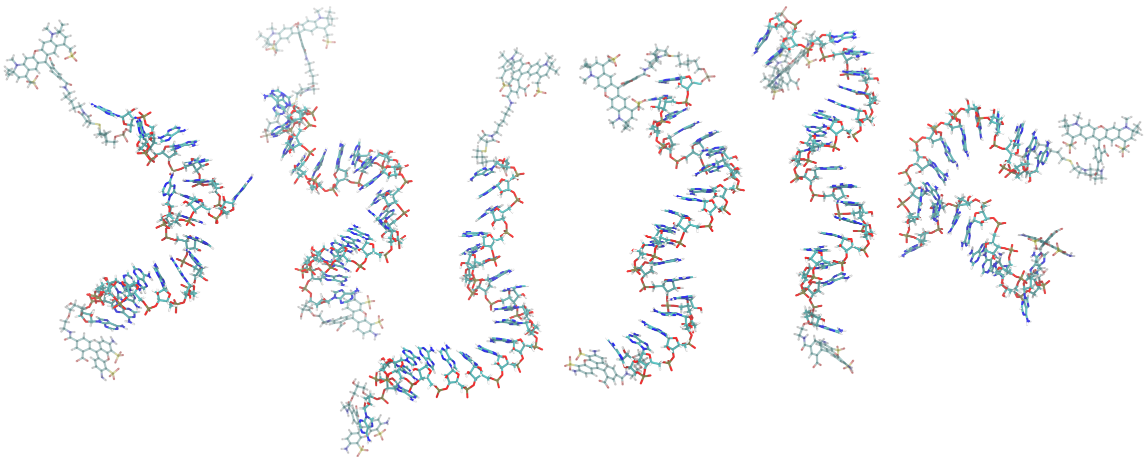

Figure S3: Snapshots of $rA_{19}$ models from HCG with attached FRET labels Alexa Fluor 594 and Alexa Fluor 488 at the 5′ and 3′ ends, respectively. MD libraries of the dye molecules were prepared previously.[S3]

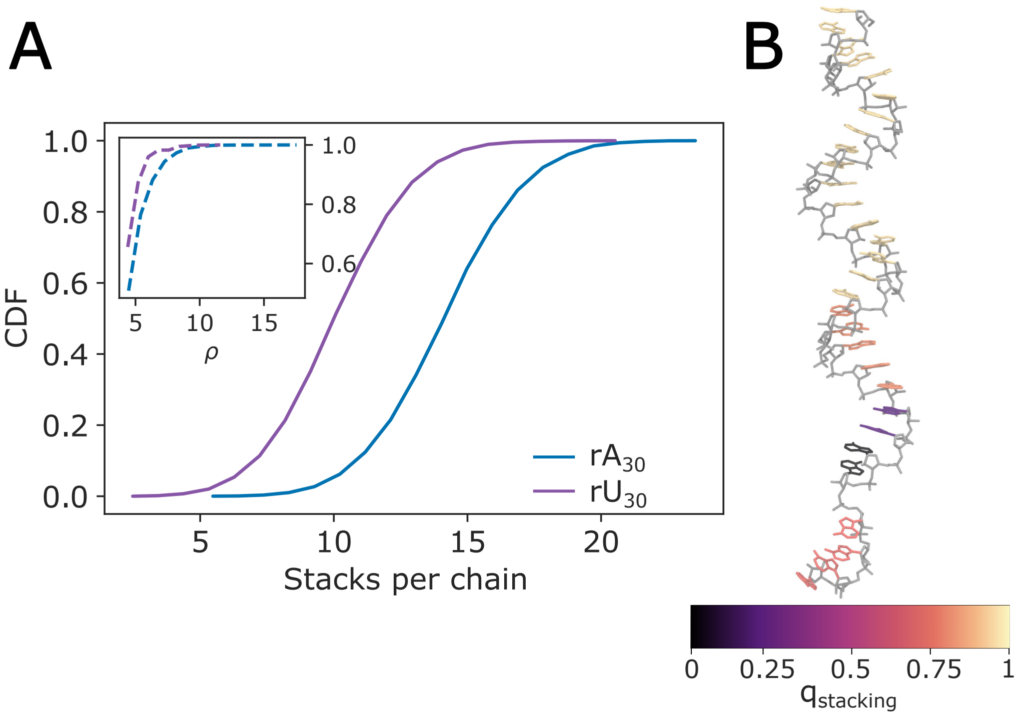


Figure S4: Stacking analysis of rA and rU 30mer ensembles grown with HCG. (A) Cumulative distribution function (CDF) of the number of stacks per chain for $rA_{30}$ in blue and $rU_{30}$ in violet. One stack is formed by two nucleobases. The inset shows the CDF of nucleobases involved in consecutive stacks ($\rho$) for both polymers. We define a consecutive stack as four or more stacked nucleobases. (B) A render of $rA_{30}$ with nucleobases coloured according to the stacking. Hydrogen atoms are omitted for clarity. The nucleic backbone and the sugar moiety are shown in gray. The stacking analysis was performed using the Barnaba package.[S1]

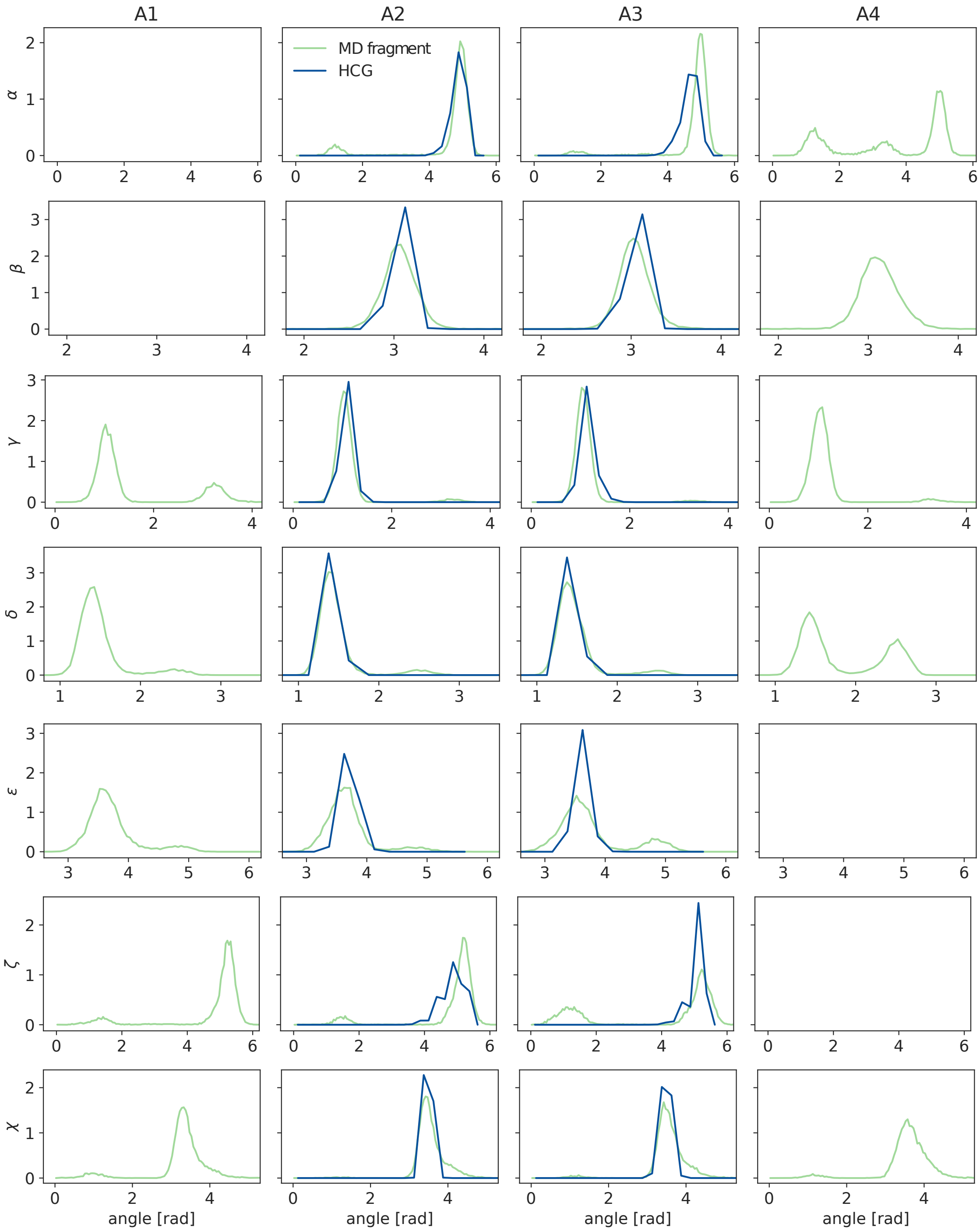


Figure S5: Distributions of backbone dihedral angles of the adenosine nucleotides as sampled in the MD fragment and the assembled $rA_{19}$ homopolymer from HCG. $\alpha$, $\beta$, $\gamma$, $\delta$, $\epsilon$, $\zeta$, and $\chi$ angle distributions of A1-A4 as sampled in the 100 ns REMD simulation of the rA tetramer fragment are shown in light green. The distributions of the backbone dihedral angles of the A2 and A3 nucleotides after being assembled into $rA_{19}$ with HCG are shown in blue. For $rA_{19}$ the average distributions of the analyzed dihedral angles across all residues that were in the second (A2) or the third (A3) position of the original $rA_4$, respectively, are shown.

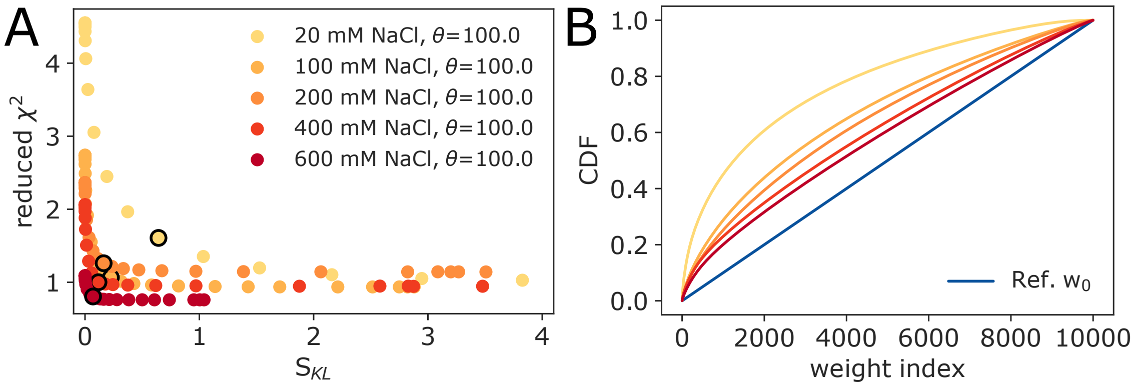


Figure S6: Assesment of the BioEn reweighting of the $rA_{30}$ structural HCG ensemble against SAXS measurements carried out at different NaCl concentrations. (A) L-curve analysis for the BioEn refinement results for NaCl concentrations 20, 100, 200, 400, and 600 mM. (B) Cumulative distribution of rank ordered weights. Uniform reference weights $w_0$ are shown in blue, refined weights from BioEn for $\theta = 100$ are shown in colors from dark red - yellow for the different salt concentrations. Color scheme as indicated in the legend of panel A.

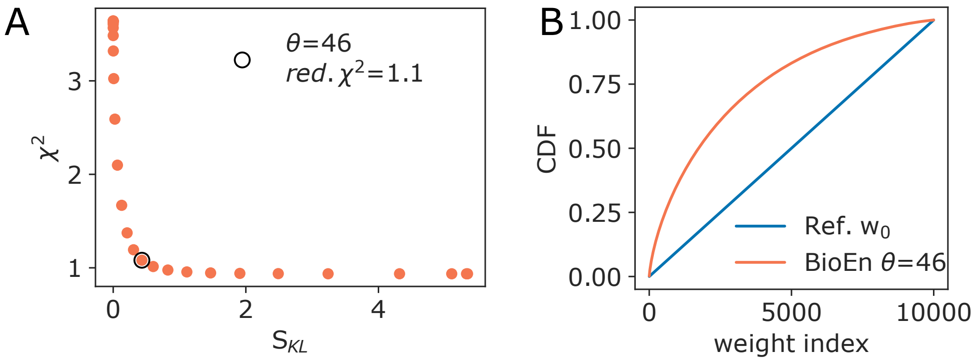


Figure S7: Ensemble refinement of a $rU_{30}$ HCG ensemble against experimental SAXS data. The experimental profile was recorded at 100 mM NaCl by Plumridge et al.[S4] (A) L-curve analysis with reduced $\chi^2$ plotted against $S_{KL}$ as a result of the BioEn reweighting. (B) Cumulative distribution of rank ordered weights for the uniform reference weights $w_0$ and refined weights for $\theta = 46$ (blue and orange, respectively).

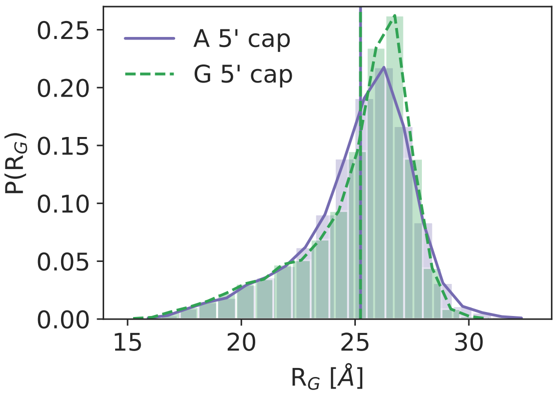


Figure S8: Distribution of the radius of gyration for HCG ensemble from homo- or heteropolymers. 10000 HCG structures were grown either from homo tetramer fragments, i.e., with adenosinemonophosphate as 5′ terminal headgroup (purple) or from hetero tetrameric fragments with guanosinemonophosphate as 5′ terminal headgroup (green, dotted lines). The range for $R_G$ measured at 100 mM and 200 mM NaCl in SAXS experiments is shown as gray shaded area.[S4]

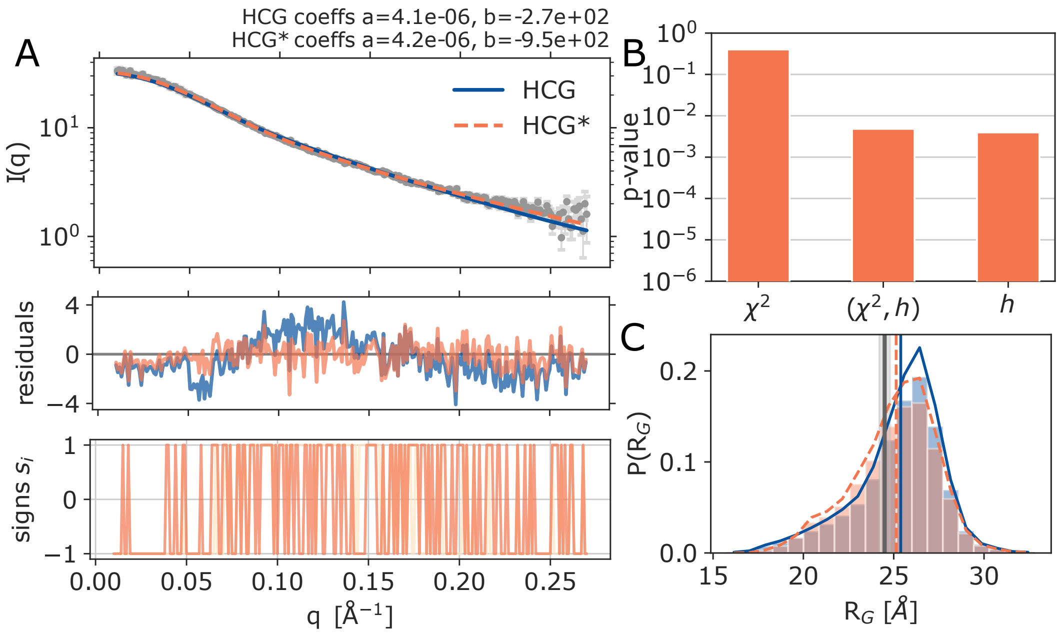


Figure S9: Ensemble refinement of a $rA_{30}$ HCG ensemble against experimental SAXS data recorded at 100 mM NaCl by Plumridge et al.[S4] (A) Top: Scattering profiles for experiment, unrefined HCG, refined HCG* ensemble (gray, blue, and orange) with the respective fitting parameters (intensity scale factor a and the background correction constant b). Middle: Residuals for the SAXS data. Bottom: Signs analysis of the residuals. (B) P-values of the refined ensemble for reduced $\chi^2$, combined reduced $\chi^2$ and h, and h as calculated by the hplusminus analysis package.[S5] (C) Unrefined and weighted distribution of $R_G$ sampled in HCG and HCG*, and their averages (blue and dotted orange). The experimental value determined at 100 mM NaCl is shown as dark gray line with the error range highlighted as light gray shaded area.

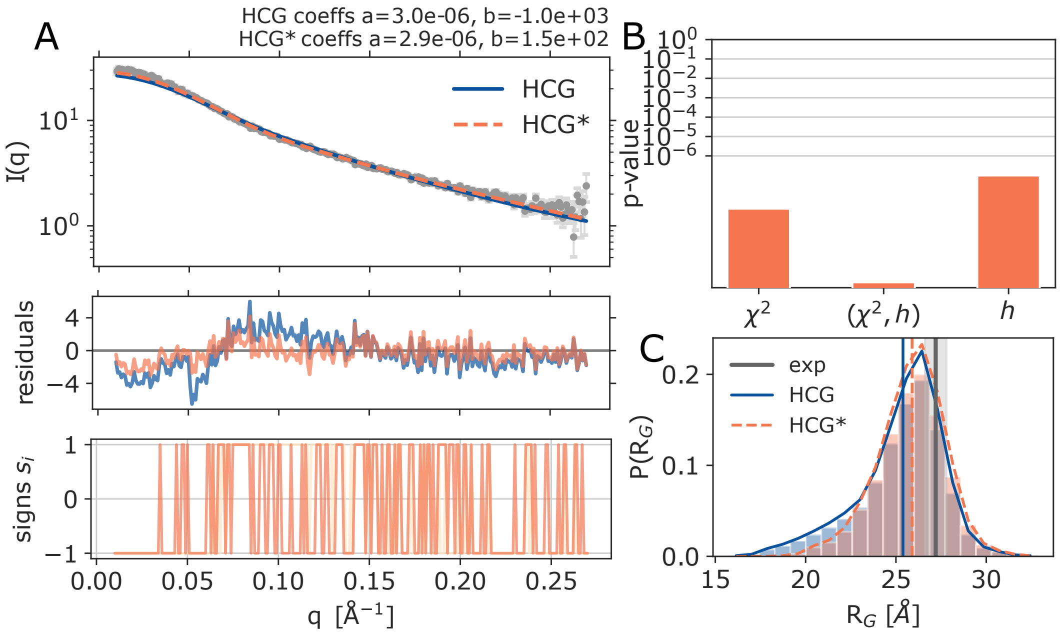


Figure S10: Ensemble refinement of a $rA_{30}$ HCG ensemble against experimental SAXS data recorded at 20 mM NaCl by Plumridge et al.[S4] (A) Top: Scattering profiles for experiment, unrefined HCG, refined HCG* ensemble (gray, blue, and orange) with the respective fitting parameters (intensity scale factor a and the background correction constant b). Middle: Residuals for the SAXS data. Bottom: Signs analysis of the residuals. (B) P-values of the refined ensemble for reduced $\chi^2$, combined reduced $\chi^2$ and h, and h as calculated by the hplusminus analysis package.[S5] (C) Unrefined and weighted distribution of $R_G$ sampled in HCG and HCG*, and their averages (blue and dotted orange). The experimental value determined at 20 mM NaCl is shown as dark gray line with the error range highlighted as light gray shaded area.

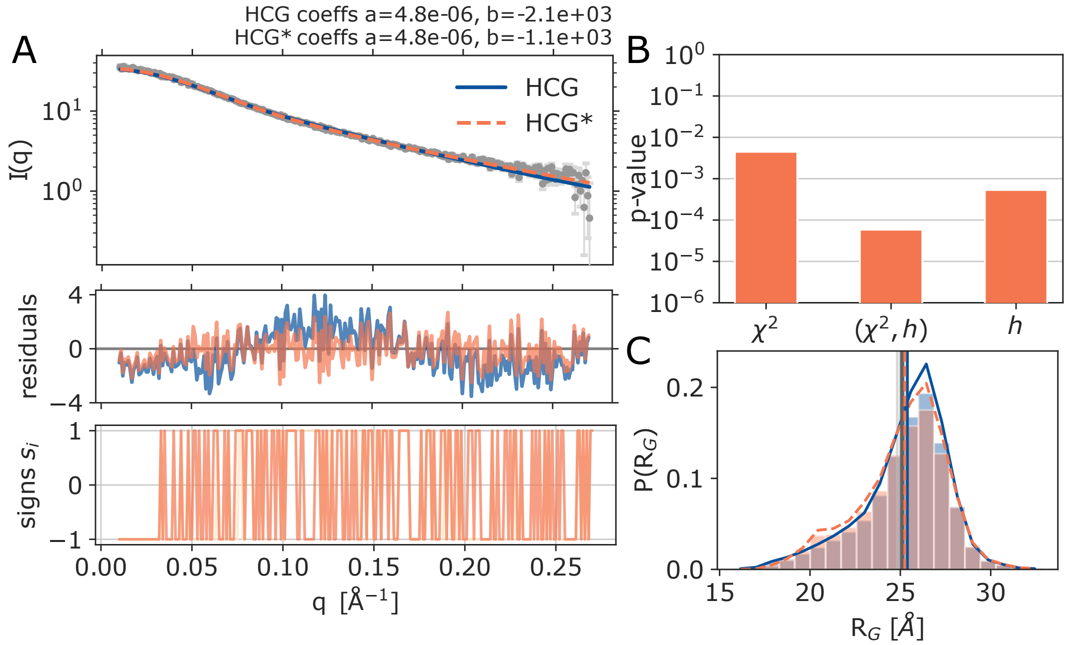


Figure S11: Ensemble refinement of a $rA_{30}$ HCG ensemble against experimental SAXS data recorded at 200 mM NaCl by Plumridge et al.[S4] (A) Top: Scattering profiles for experiment, unrefined HCG, refined HCG* ensemble (gray, blue, and orange) with the respective fitting parameters (intensity scale factor a and the background correction constant b). Middle: Residuals for the SAXS data. Bottom: Signs analysis of the residuals. (B) P-values of the refined ensemble for reduced $\chi^2$, combined reduced $\chi^2$ and h, and h as calculated by the hplusminus analysis package.[S5] (C) Unrefined and weighted distribution of $R_G$ sampled in HCG and HCG*, and their averages (blue and dotted orange). The experimental value determined at 200 mM NaCl is shown as dark gray line with the error range highlighted as light gray shaded area.

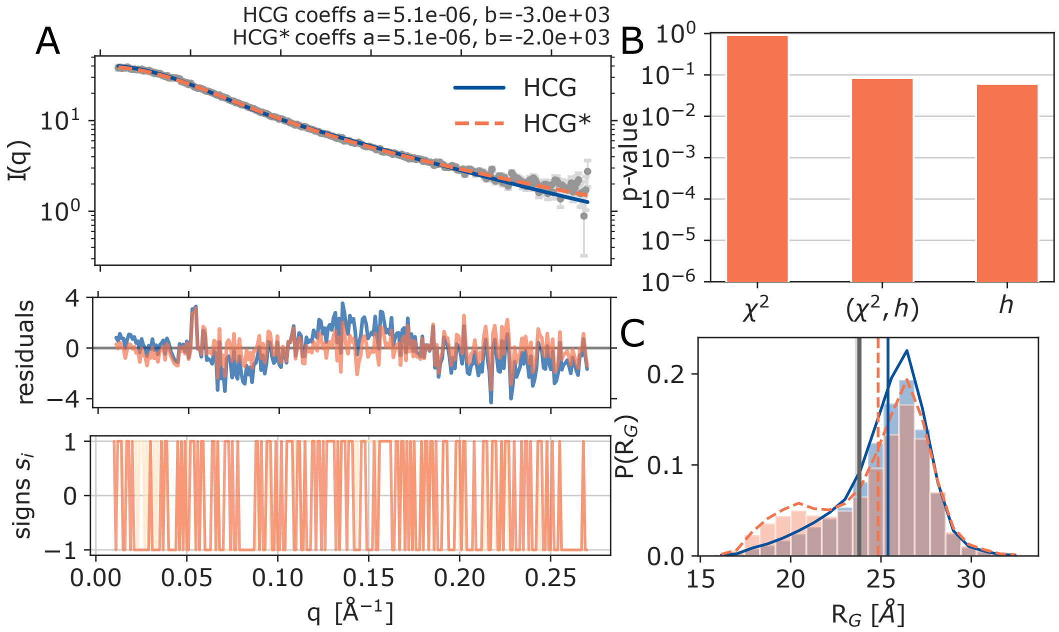


Figure S12: Ensemble refinement of a $rA_{30}$ HCG ensemble against experimental SAXS data recorded at 400 mM NaCl by Plumridge et al.[S4] (A) Top: Scattering profiles for experiment, unrefined HCG, refined HCG* ensemble (gray, blue, and orange) with the respective fitting parameters (intensity scale factor a and the background correction constant b). Middle: Residuals for the SAXS data. Bottom: Signs analysis of the residuals. (B) P-values of the refined ensemble for reduced $\chi^2$, combined reduced $\chi^2$ and h, and h as calculated by the hplusminus analysis package.[S5] (C) Unrefined and weighted distribution of $R_G$ sampled in HCG and HCG*, and their averages (blue and dotted orange). The experimental value determined at 400 mM NaCl is shown as dark gray line with the error range highlighted as light gray shaded area.

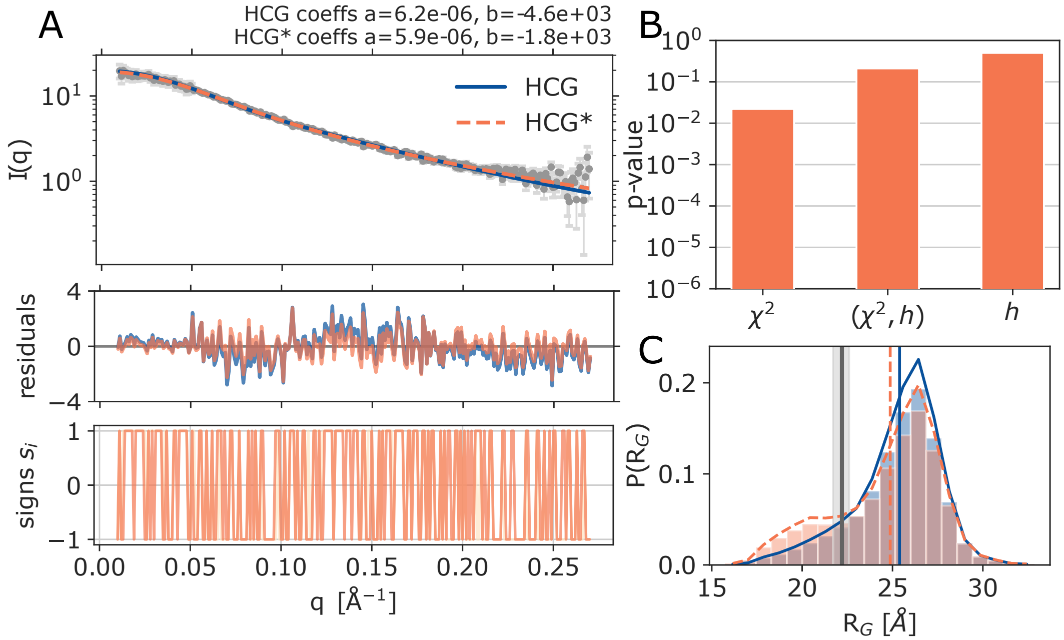


Figure S13: Ensemble refinement of a $rA_{30}$ HCG ensemble against experimental SAXS data recorded at 600 mM NaCl by Plumridge et al.[S4] (A) Top: Scattering profiles for experiment, unrefined HCG, refined HCG* ensemble (gray, blue, and orange) with the respective fitting parameters (intensity scale factor a and the background correction constant b). Middle: Residuals for the SAXS data. Bottom: Signs analysis of the residuals. (B) P-values of the refined ensemble for reduced $\chi^2$, combined reduced $\chi^2$ and h, and h as calculated by the hplusminus analysis package.[S5] (C) Unrefined and weighted distribution of $R_G$ sampled in HCG and HCG*, and their averages (blue and dotted orange). The experimental value determined at 600 mM NaCl is shown as dark gray line with the error range highlighted as light gray shaded area.

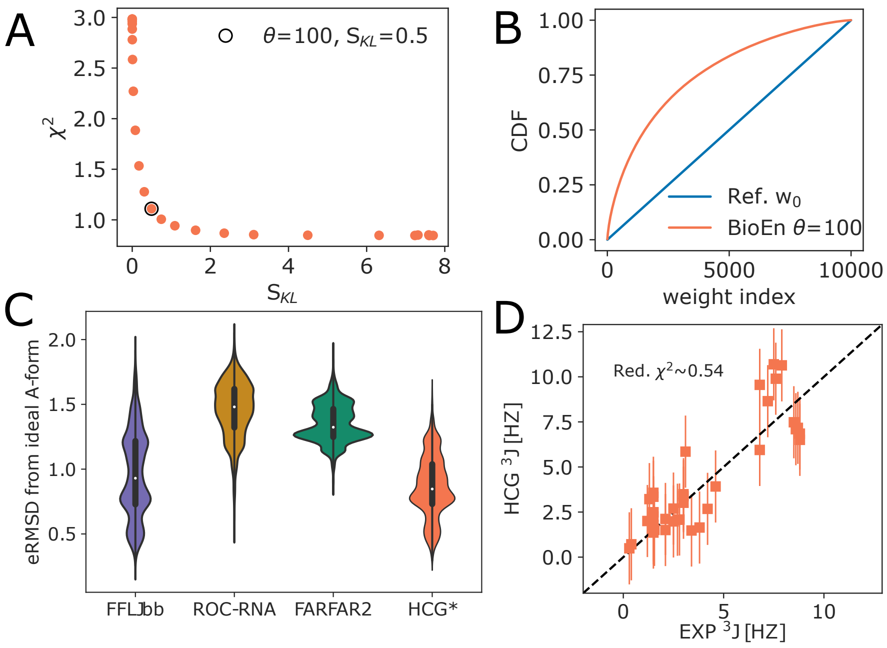


Figure S14: Structural ensembles of heteropolymeric UCAAUC from HCG reweighted against the experimental scattering profile.[S6] (A) L-curve analysis of the BioEn refinement. We chose refined weights for $\theta = 100$ for further analysis of properties shown here. (B) Cumulative distribution of rank-ordered weights. Uniform reference weights are shown in blue, refined weights for $\theta = 100$ in orange. (C) Distribution of the eRMSD to ideal A-form. (D) Correlation plot of $^3$J-couplings of experimentally measured[S7] and calculated values for the refined HCG* ensemble. Error estimates for HCG* of 2 Hz are indicated as vertical lines.

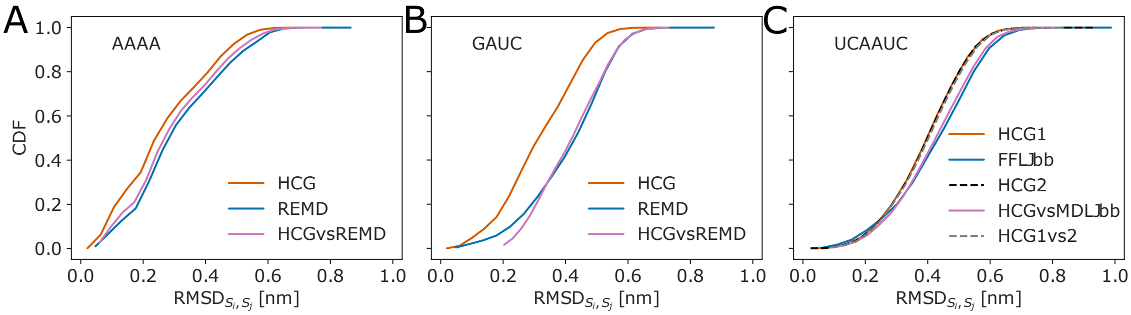


Figure S15: Cumulative distribution of pairwise RMSDs in (RE)MD ensembles and HCG ensembles of ssRNA polymers with different lengths. CDFs are shown for the pairwise RMSD within (RE)MD simulations (blue), within HCG ensembles (dark orange and dotted black), and between random structures S$i$ and S$j$ in different ensembles (rose and gray). Pairwise RMSDs were calculated using all heavy atoms within a polymer. Each polymer ensemble sampled with either (RE)MD or HCG contained 10000 structures. (A) AAAA and (B) GAUC tetramer, REMD run for 100 ns with the DESRES force field combined with the TIP4P-d water model. (C) UCAAUC hexamer with the MD trajectory run for 1 $\mu$s with the LJbb force field by Bergonzo et al.[S6] Here, the RMSD between two independent HCG ensembles of UCAAUC is also shown (dotted gray).

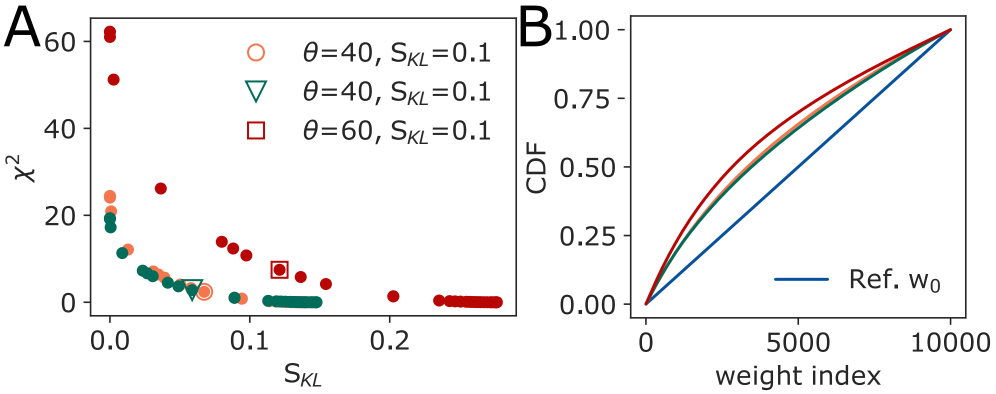


Figure S16: Assessment of the refinement of the structural HCG ensemble of $rA_{19}$ against single-molecule FRET experiments. We reweighted the distribution of FRET efficiencies calculated from a HCG ensemble with 10000 chains of $rA_{19}$ with mapped dyes calculted with model 1 (dynamic dye orientations, orange), model 2 (dynamic dyes, dark green), or model 3 (static dyes,[S8] dark red) against the experimental mean efficiency <E> using BioEn.[S9] (A) L-curve analysis of the results from reweighting. We found $S_{KL} < 0.1$ and $\chi^2 \approx 2.7$ for a set of weights for $\theta = 40$ for model 1 (orange empty circle). For model 2 we chose weights for $\theta = 40$ with $S_{KL} < 0.1$ and $\chi^2 \approx 7.8$ (dark green empty triangle). For model 3 we chose a set of weights for $\theta = 60$ with $S_{KL} \approx 1.4$ and $\chi^2 \approx 8$ (dark red empty square). (B) Cumulative distribution of rank ordered weights for the respective $\theta$. Color scheme as indicated in panel A.

### References


(S1) Bottaro, S.; Bussi, G.; Pinamonti, G.; Reiber, S.; Boomsma, W.; Lindorff-Larsen, K. Barnaba: software for analysis of nucleic acid structures and trajectories. *RNA* **2019**, *25*, 219–231.

(S2) Bottaro, S.; Bussi, G.; Lindorff-Larsen, K. Conformational ensembles of noncoding elements in the SARS-CoV-2 genome from molecular dynamics simulations. *J. Am. Chem. Soc.* **2021**, *143*, 8333–8343.

(S3) Grotz, K. K.; Nuuesch, M. F.; Holmstrom, E. D.; Heinz, M.; Stelzl, L. S.; Schuler, B.; Hummer, G. Dispersion correction alleviates dye stacking of single-stranded DNA and RNA in simulations of single-molecule fluorescence experiments. *J. Phys. Chem. B* **2018**, *122*, 11626–11639.

(S4) Plumridge, A.; Andresen, K.; Pollack, L. Visualizing disordered single-stranded RNA: connecting sequence, structure, and electrostatics. *J. Am. Chem. Soc.* **2020**, *142*, 109–119.

(S5) Köfinger, J.; Hummer, G.; Köfinger, J. Powerful statistical tests for ordered data. *ChemRxiv* **2021**,

(S6) Bergonzo, C.; Grishaev, A.; Bottaro, S. Conformational heterogeneity of UCAAUC RNA oligonucleotide from molecular dynamics simulations, SAXS, and NMR experiments. *RNA* **2022**, *28*, 937–946.

(S7) Zhao, J.; Kennedy, S. D.; Berger, K. D.; Turner, D. H. Nuclear magnetic resonance of single-stranded RNAs and DNAs of CAAU and UCAAUC as benchmarks for molecular dynamics simulations. *J. Chem. Theory Comput.* **2020**, *16*, 1968–1984.

(S8) Hummer, G.; Szabo, A. Dynamics of the orientational factor in fluorescence resonance energy transfer. *J. Phys. Chem. B* **2017**, *121*, 3331–3339.

(S9) Hummer, G.; Köfinger, J. Bayesian ensemble refinement by replica simulations and reweighting. *J. Chem. Phys.* **2015**, *143*.
